# Is DNA metabarcoding an option for formaldehyde-preserved zooplankton time series?

**DOI:** 10.64898/2026.02.06.704415

**Authors:** Aitor Albaina, Anders Lanzén, Irati Miguel, Fernando Rendo, María Santos

## Abstract

The recovery of amplifiable DNA from formaldehyde□fixed (FF) zooplankton samples has long been considered unfeasible. Nevertheless, advances in DNA sequencing and methods for retrieving highly degraded genetic material have demonstrated that even million□year□old samples and FF museum specimens can yield usable DNA. To access the biological information preserved in long□term zooplankton time series, we assessed methodologies for extracting amplifiable DNA from community samples stored for up to 28 years in formaldehyde at room temperature.

On one hand, we report the failure of a method previously described as successful for FF zooplankton samples, likely due to the cold□storage conditions (4□°C) used in the original study. On the other hand, by adapting two extraction protocols designed for FF museum specimens—representing harsher and softer alternatives (HHA and HPC, respectively)—we successfully amplified and sequenced a subset of FF zooplankton samples. As expected, DNA integrity and sample pH were inversely related to preservation time, and only short DNA fragments were recovered, ruling out the use of commonly employed ≥300□bp metabarcoding markers. While DNA integrity appeared to be a better predictor than DNA yield for amplification success, the presence of a gel band of the expected size did not always guarantee congruence with microscopy□based assessments. Although amplifiable DNA was recovered from most samples, including some of the oldest, community compositions concordant with microscopy were consistently recovered only from samples preserved for up to two years. Beyond this point, the HHA and HPC methods produced divergent results, reflecting a trade□off between the removal of formaldehyde□induced cross□linkages and the avoidance of additional DNA damage. Among the small universal markers tested (∼120–170□bp), including one nuclear rRNA marker and two mitochondrial markers, only the 18S rRNA V9 region consistently amplified. We conclude by providing a set of recommendations aimed at improving the methods presented here.

## INTRODUCTION

Traditionally, zooplankton has been fixed and preserved in buffered formaldehyde (usually 4% formaldehyde; sometimes referred as 10% formalin; see Simmons 2014 for a review) due to its low price and, mainly, to its optimal performance preserving morphological features in the long term (e.g. UNESCO, 1976; Mauchline, 1998; ICES Zooplankton Methodology Manual 2000; Bucklin and Allen, 2004; Suthers et al., 2019). This applies to the vast majority of zooplankton long time series that has been archiving zooplankton samples regularly since the first half of 20^th^ century (e.g. since 1931 and 1951 for the pioneers Continuous Plankton Recorder, CPR, and California Cooperative Oceanic Fisheries Investigations, CalCOFI, respectively; see https://www.st.nmfs.noaa.gov/copepod/time-series/ and ÓBrien et al., 2017, for a compilation of zooplankton time series). Zooplankton communities monitoring has, since then, proven to be of huge utility when assessing water quality and, more recently, the response of the aquatic realm to the ongoing climate change (e.g. Edwards et al., 2010; Holland et al., 2023, 2025; Ratnarajah et al., 2023; Chust et al., 2024; Grigoratou et al., 2025).

Until the advent of high-throughput image and DNA-bases approaches, zooplankton identification relayed on expert taxonomist inspection under the microscope, a highly costly, both in time and money, approach requiring a deep training with a pronounced learning curve. Moreover, due to the inherent difficulty of zooplankton species discrimination, the bulk of zooplankton taxonomists is specialized in one or a few animal groups (e.g. copepods or cnidarians) (Peters et al., 2025). Because of these limitations and with the aim of generalizing the use of zooplankton monitoring to inform policy makers, in the last decades the use of alternative methods has been fostered. While image-based approaches (e.g. ZooScan, Underwater Video Profiler, ZooCAM, FlowCam; see e.g. Eerola et al., 2024 for a review) seemed promising when characterizing zooplankton communities in low diversity systems such as fjords (e.g. Naito et al., 2019), they are still far from the taxonomic resolution of skilled taxonomists and are not expected to ever replace them (e.g. Cornils et al., 2022; Ershova et al., 2025). However, they have revealed as a clear alternative to estimate size-structured biomasses. More recently, DNA metabarcoding, where one or, ideally, several “universal” barcodes, are sequenced from whole community/net samples, following Polymerase Chain Reaction (PCR)-based amplification, allowing a species level taxonomy even in cryptic groups (providing they are covered by the reference database) has gained momentum. This is due to the decreasing cost of DNA sequencing and the growing population of reference databases for zooplankton (e.g. Bucklin et al., 2016; Laakmann et al., 2020; Bucklin et al., 2021). In this sense, DNA metabarcoding of zooplankton communities usually retrieves a higher number of taxa than microscopy by an expert taxonomist (e.g. Keck et al., 2022). Nowadays, the combination of 1) several complementary barcodes (typically, a Cytochrome Oxidase I –“COI”- gene fragment and a ribosomal RNA -rRNA- one; e.g. Albaina et al. 2024 for a review) with 2) parallel mock samples analysis, as to correct for potential biases (e.g. Santoferrara, 2019), offers the promise of a semi-quantitative method with unrivalled taxonomic resolution, at least for the bulk of taxa in a zooplankton sample (e.g. Shelton et al., 2023).

However, formaldehyde preservation represents a significant impediment to DNA amplification and sequencing, which worsens with increased fixation time (e.g. Bucklin and Allen, 2004; Bucklin et al., 2011; Gilbert et al., 2007; Straube et al., 2021; Hahn et al., 2022; Vezzulli et al., 2022; Steiert et al., 2023). This is due to three factors:

1. Formaldehyde decreases solutiońs pH due to its gradual degeneration to formic acid (e.g. Hahn et al., 2022), therefore involving DNA fragmentation that potentially will impede even short length barcode amplification and thus sequencing (e.g. Díaz-Viloria et al. 2005). This impact is to be at least partly delayed if a) formaldehyde is buffered (e.g. Miething et al., 2006; Steiert et al., 2023), b) the pH is regularly controlled as to maintain the buffer effect and c) the sample if storage in cold (4° C) conditions (Koshiba et al., 1993). While the first countermeasure is nowadays generalized for zooplankton samples (usually with borax, di-Sodium tetra-Borate 10-hydrate, and to a pH of ∼8; e.g. ICES Zooplankton Methodology Manual 2000), the other two measures, and especially the latter, are not (especially in the long time series due to obvious logistic constraints).
2. Formaldehyde promotes cross-linking within the DNA itself or between DNA and proteins (histones) impeding the physical access of PCR polymerases to the DNA template and, therefore, preventing/inhibiting DNA (barcode) amplification (e.g. Hoffman et al., 2015; Oka et al., 2024). And,
3. Formaldehyde is capable of producing DNÁs base modifications, therefore potentially creating a false negative/positive presence signal (e.g. De Giorgi et al., 1994; Quach et al., 2004)

Consequently, formaldehyde□preserved zooplankton samples have traditionally been regarded as unsuitable for DNA□based methodologies.

However, the pioneer work of Bucklin and Allen (2004) already reported that, if DNA extraction methods are adapted to overcome the damages associated to formaldehyde fixation, short DNA fragments (< 200 bp) can be amplified from zooplankton specimens (krill individuals stored up to 25 years in formaldehyde), albeit with a very low success rate (but see Ripley et al., 2008 that failed when applying the same method for a 100 bp amplicon, and even after just one week in formaldehyde).

Since then, DNA extraction methods and sequencing technologies have experienced tremendous advances, allowing for example the amplification of ancient DNA fragments (> 1M years old) with DNA metabarcoding approaches (Kjær et al., 2022) or the assembly of the Neanderthal genome. Regarding formaldehyde-fixed (FF) tissues, although the fragmentation of DNA is not amendable (making the use of short length barcodes mandatory; e.g. Gilbert et al., 2007), recent improvements in DNA extraction methods, mainly developed to access DNA from human tissués Formaldehyde-Fixed Paraffin-Embedded (FFPE) samples, but also later applied to museumś animal specimens, has proven successfully to counteract, at least partly, the DNA-protein cross-links and the artificial base modifications (e.g. Holleley and Hahn, 2025). More in detail, the use of both high temperatures and pH at a certain point when digesting the tissue sample (the so-called “hot-alkaline lysis”; from now on ‘HÁ approach) has been shown effective to remove cross-links, allowing target DNA amplification (Campos and Gilbert, 2012, Straube et al., 2021; Steiert et al., 2023, Hahn et al., 2022, 2024a). However, this comes with a trade-off as these harsh conditions can further damage DNA (e.g. Totoiu et al., 2020). A “softer” alternative increases the sample lysis/digestion step with proteinase-K (pK) for up to several days (e.g. Ripley et al., 2008; Palero et al., 2010) as pK activity can also remove cross-links (e.g. Totoiu et al., 2020; Straube et al., 2021). Apart from this, formaldehyde-caused base modifications can be, at least partly, corrected with *ad hoc* designed enzyme cocktails such as the NEBNext® FFPE DNA Repair Mix one (New England Biolabs). However, these repair kits were originally designed for human FFPE tissue samples and hardly applied to other sample matrices (Gilbert et al. 2007 and Steiert et al., 2023 for a review; but see e.g. Hahn et al., 2022 and Shiozaki et al. 2021 for application with museum animal specimens and zooplankton samples, respectively).

Within this context, in the last years, different authors have reported a relatively success when amplifying DNA from formaldehyde-preserved plankton samples. On the one side, Shiozaki et al. (2021) reported the amplification of a relatively long barcode (the rRNA 18S V7-V8 one, with 450 bp length) in zooplankton community samples with formaldehyde preservation times of up to 23 years; however, their samples had been maintained at 4°C since collection, a condition not frequent with archived zooplankton samples. As far as we know, no other work has tried DNA metabarcoding of formaldehyde-fixed zooplankton community samples; the rest of works facing DNA amplification of individual specimens by means of barcoding with Sanger sequencing. On the other side, Vezzulli et al. (2012, 2022), were able to amplify a short DNA fragment (113 bp length) from CPR plankton community samples, preserved at RT conditions, from up to 50 years. However, although promising due to the long preservation times, these authors did not apply DNA metabarcoding, but a species-specific real time PCR approach, and, more importantly, neither targeted Metazoan DNA but a bacterial DNA fragment (*Vibrio* spp.).

With the aim of getting access to the priceless genetic treasure of formaldehyde-preserved archived zooplankton samples, we tested these promising methodologies with samples covering a wide range of preservation times (from two to twenty-eight years). For this, we selected samples from the BIOMAN (BIOMass of ANchovy) surveys, an annual scientific campaign to estimate the biomass of European anchovy (*Engraulis encrasicolus*) in the Bay of Biscay (BoB) following the Daily Egg Production Method (DEPM). BIOMAN surveys have been conducted annually since 1987 (except for 1993), covering the anchovýs peak reproduction period (from the beginning to the last quarter of May). More in detail, zooplankton samples, where anchovieś eggs are sorted out for the DEPM, are collected in an adaptive survey covering the whole anchovýs reproduction area in a high spatial resolution grid of 3*15 nautical miles encompassing the BoB coastal, shelf and shelf-break ecosystems (e.g. Albaina et al. 2007b, Santos et al. 2023). As a results of this, an average of ∼500 mesozooplankton samples (150 μm mesh net) are archived annually in the BIOMAN plankton collection. While a small fraction of the time series has been characterized under the microscope pointing to a high diverse and mesoscale-structured zooplankton community (Albaina et al., 2004, 2007a, 2007b), the rest remain in 4% buffered formaldehyde and RT conditions. This represents a huge archive covering the last 40 yearś spring period in a transitional region between the tropical and the temperate-boreal realms that could shed light on the impact of climate change in the base of marine communities.

Aiming to retrieve the maximum information from this valuable long time series and, at the same time, develop a methodology scalable to other comparable zooplankton time series worldwide, we herein adapted and applied three novel and promising DNA extraction methods to a set of BIOMAN zooplankton samples covering the bulk of the serieś temporal period. The attained results and recommendations are to be considered as a starting point aiming to develop a highly successful methodology to recover DNA information from archived zooplankton samples by means of DNA metabarcoding.

## MATERIAL AND METHODS

### Zooplankton sampling

A total of 22 zooplankton samples collected in the Basque Country surveys to estimate the biomass of anchovy (*Engraulis encrasicolus*) (BIOMAN campaigns, e.g. Albaina et al. 2007) and preserved in formaldehyde were selected for the present study. Samples were collected using vertical hauls of a 150 μm PAIROVET net fitted with a flowmeter and lowered to a maximum depth of either 100 or 5 m above the bottom in shallower stations (e.g. Albaina et al. 2004).Net samples were preserved immediately after collection with 4% borax buffered formaldehyde (to a pH of 7.5-8) and stored at room temperature (RT) since then. Five samples along a cross-shelf transect in front of the mouth of the Gironde estuary (45.62° N, and from -1.50 to -3.35°W), encompassing one coastal, two inner shelf and two outer shelf stations (C, IS1, IS2, OS1 and OS2, respectively, with depths of 39, 71, 76, 138 and 149 m; Figure 1) were selected from four different years (1995, 2004, 2011 and 2017) as to cover a range of up to 28 years in formaldehyde (Table 1). Apart from this, two coastal locations (crosses in Figure 1, with depths of 30 and 55 m) sampled in 2022, with a lower fixation time (two years; samples C1 and C2) were also included for comparison. Finally, a second consecutive PAIROVET haul performed in the very same 2022 locations was preserved immediately after collection in 100% ethanol for positive controls (C+1, C+2).

**Figure 1.**
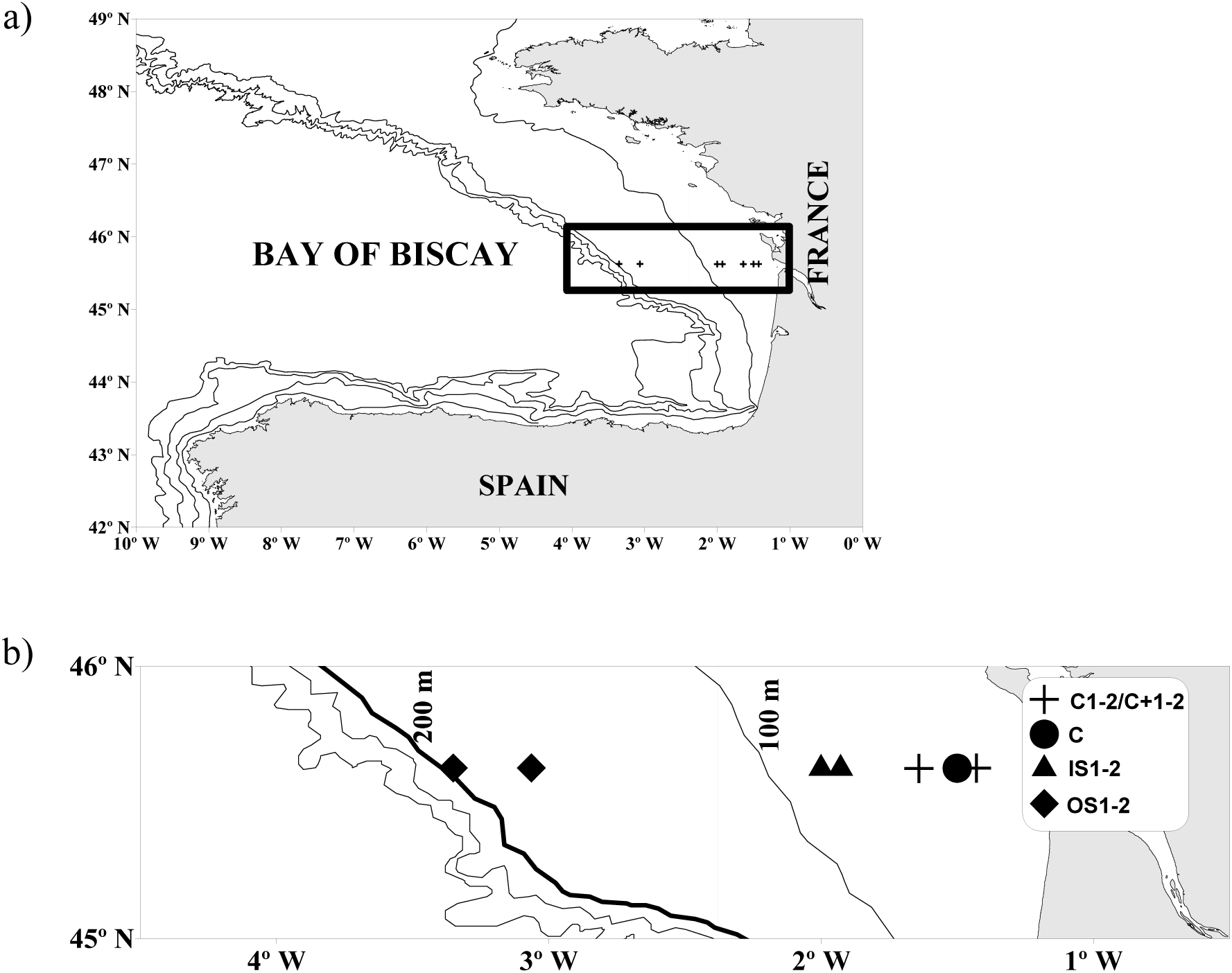
Stationś map. Five stations along a cross-shelf transect in the Bay of Biscay were sampled at four different years in the BIOMAN campaigns (1995, 2004, 2011 and 2017); these corresponded to costal (C; circle symbol), internal shelf (IS1 and IS2; up-triangle) and outer shelf samples (OS1 and OS2; diamond). In 2022, two costal samples (crosses) were preserved both in ethanol (C+1, C+2) and in formaldehyde (C1, C2) for positive control and comparison against a lower time in formaldehyde, respectively.

**Table 1.** Sample details. Sampling details including BIOMAN campaign, coordinates and bottom depth are shown, along with the pH and preservation time in formaldehyde before DNA extraction along with sea surface temperature and salinity (ST and SS, respectively. The two positive controls fixed in ethanol, 22/C+1, 22/C+2, are included (total n=24). Sampleś CODEs are defined based in the sampling year and the location along the sampled transect (see Figure 1). Values of ≤ 5.5 pH units correspond to the lower limit of detection (pH test-strips). NA: not available

| YEAR | CODE | TIME IN FORMALIN | pH | LATITUDE | LONGITUDE | DEPTH | ST | SS |
| --- | --- | --- | --- | --- | --- | --- | --- | --- |
| 1995 | **95/C** | 28 years | ≤ 5.5 | 45.63 | -1.50 | 40 | 13.6 | 34.3 |
| 1995 | **95/IS1** | 28 years | ≤ 5.5 | 45.63 | -1.93 | 70 | 13.7 | 34.5 |
| 1995 | **95/IS2** | 28 years | ≤ 5.5 | 45.62 | -2.00 | 76 | 13.8 | NA |
| 1995 | **95/OS1** | 28 years | ≤ 5.5 | 45.62 | -3.28 | 138 | 14.1 | NA |
| 1995 | **95/OS2** | 28 years | ≤ 5.5 | 45.62 | -3.35 | 147 | 14.0 | NA |
| 2004 | **04/C** | 18 years | ≤ 5.5 | 45.62 | -1.50 | 39 | 14.2 | 28.9 |
| 2004 | **04/IS1** | 18 years | ≤ 5.5 | 45.62 | -1.93 | 72 | 14.2 | 33.1 |
| 2004 | **04/IS2** | 18 years | ≤ 5.5 | 45.62 | -2.00 | 76 | 14.0 | 33.7 |
| 2004 | **04/OS1** | 18 years | ≤ 5.5 | 45.62 | -3.28 | 139 | 13.3 | 35.3 |
| 2004 | **04/OS2** | 18 years | ≤ 5.5 | 45.62 | -3.35 | 151 | 13.2 | 35.3 |
| 2011 | **11/C** | 12 years | 6 | 45.62 | -1.50 | 38 | 15.1 | 34.5 |
| 2011 | **11/IS1** | 12 years | 6 | 45.62 | -1.93 | 70 | 17.4 | 34.6 |
| 2011 | **11/IS2** | 12 years | 6 | 45.63 | -2.00 | 75 | 17.2 | 34.6 |
| 2011 | **11/OS1** | 12 years | 6 | 45.63 | -3.28 | 137 | 16.8 | 35.7 |
| 2011 | **11/OS2** | 12 years | 6 | 45.63 | -3.35 | 149 | 16.8 | 35.7 |
| 2017 | **17/C** | 7 years | 6.5 | 45.63 | -1.50 | 38 | 15.0 | 33.0 |
| 2017 | **17/IS1** | 7 years | 6.5 | 45.62 | -1.93 | 72 | 14.9 | 34.4 |
| 2017 | **17/IS2** | 7 years | 6.5 | 45.62 | -2.00 | 78 | 14.8 | 34.5 |
| 2017 | **17/OS1** | 7 years | 6.5 | 45.63 | -3.06 | 137 | 14.5 | 35.6 |
| 2017 | **17/OS2** | 7 years | 6.5 | 45.63 | -3.35 | 151 | 14.3 | 35.6 |
| 2022 | **22/C1** | 2 years | 7.5 | 45.63 | -1.43 | 30 | 15.4 | 32.6 |
| 2022 | **22/C2** | 2 years | 7.5 | 45.62 | -1.64 | 55 | 18.7 | 33.6 |
| 2022 | **22/C+1** | NO | EtOH | 45.63 | -1.43 | 30 | 15.4 | 32.6 |
| 2022 | **22/C+2** | NO | EtOH | 45.62 | -1.64 | 55 | 18.7 | 33.6 |

### Microscopy characterization

Zooplankton characterization was carried out under a stereoscopic microscope. Samples from 1995 and 2004 had been already identified under the stereomicroscope (Albaina and Irigoien 2004 and 2007, respectively); the remaining samples were identified following the same procedure. More in detail, identification was made to species or genus level in the majority of the holoplanktonic groups, and to general categories in meroplanktonic forms (Table 2). In each sample, a minimum of 200 individuals (all categories included) were counted. Note that the calanoid copepod *Calanoides carinatus* is now correctly named as *Calanoides natalis* (Bradford-Grieve et al., 2017). Copepod biomasses (mg C dry weight m^−3^) were estimated using available length-biomass regressions and assuming a 40% of carbon content in total dry weight (Bamstedt 1986) as in Albaina and Irigoien (2004).

**Table 2.**
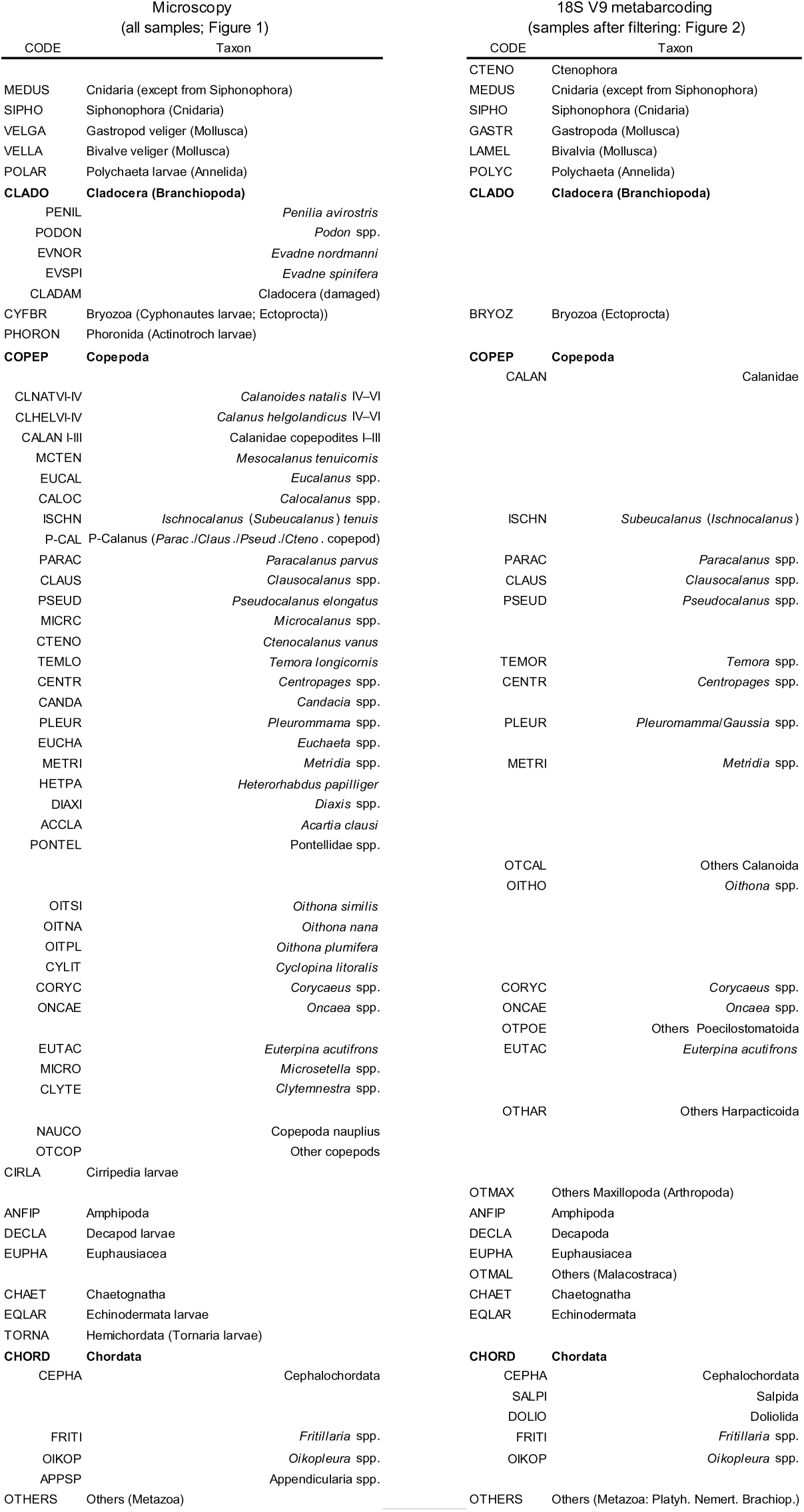
Taxonomic list. Taxa identified by microscopy (left) and 18S V9 metabarcoding (right) taxa. CODEs for graphic representations. Only taxa in those samples fulfilling bioinformatic filtering thresholds (Figure 2) are shown for 18S V9 metabarcoding.

### DNA extraction, library preparation and metabarcoding

DNA extraction, library preparation and metabarcoding were performed in the Sequencing and Genotyping Unit of the Genomic Facility by the University of the Basque Country (SGIKER, UPV/EHU). Two sequencing efforts (runs #1 and #2) were carried out for this test. For the first one, 1995, 2004 and 2011 formaldehyde-fixed (FF) samples were extracted in the summer of 2022 along with the 2022 positive controls. Due to the low PCR amplification success, more recent FF samples, i.e. 2017 and 2022 ones were extracted in 2024 for a second run (Table 1). All zooplankton samples were transferred to 96 % ethanol (EtOH) immediately before being sent to the sequencing facility due to safety requirements of the UPV/EHU. Just before, the pH of the original FF sample was measured with a pH test-strip (sensitivity range: 5.5-9) and recorded (Table 1). At the SGIKER, ethanol was removed before beginning the DNA extraction protocol.

Formaldehyde-fixed samples have been shown to be particularly prone to exogenous DNA contamination (e.g. Bucklin and Allen, 2004), a key factor to control with every damaged DNA source (e.g. Drake et al., 2022). To avoid contamination, disposable filter tips were used in every step, and lab surfaces were decontaminated after each DNA extraction session. Pre- and post-PCR areas were physically separate, with a high-pressure doors system, and with dedicated equipment and workflows. Moreover, negative controls were included in the extraction (“extraction blanks”) and in both the library preparation and sequencing (non-template controls; “NTC blanks”) steps.

Based in previous works reporting successful amplifications with some long-standing FF animal samples, including one originally designed for zooplankton, we herein tested three extraction methods: we adapted two protocols from Hahn et al. (2022), successfully applied to museum specimens, and replicated the one designed by Shiozaki et al. (2021) for zooplankton community samples. Full protocolś details are available in the Supplementary Material. The three methods represent a gradation from a “soft” method (a digestion step with pH of 8 and a 24h incubation at 65°C), that can be automatable, to a “harsh” method with very high both pH and temperatures in the lysis stage (13 and 100°C for 40 minutes, respectively). The intermediate one (the “Shiozaki” one; SH from now onwards) comprises a digestion step with pH of 11 and 24h at 65°C. More in detail, every extraction method, including the commercial kit one used for the positive controls, began with a double round of sample homogenization with Precellys®. Initially, the plankton pellet was subjected to dry homogenization, followed by a second homogenization in the lysis/digestion buffer specific to each of the extraction methods, prior to proceeding with the subsequent method-specific steps. Between 150 and 250 mg of the homogenized plankton pellet were used for each extraction method.

Briefly, on the one side, a lysis step combining a buffer of pH of 12-13 with a heat shock of 40 minutes at 100°C, without a pK digestion step, and followed by a phenol-chloroform extraction is the approach of our so-called “HHA” method (from “Hahn-Hot-Alkaline”) adapted from Hahn et al. (2022, 2024a) and Campos and Gilbert (2012). On the other side, a digestion step with pK in lower but still alkaline pH buffer conditions and without a heat shock corresponds to the “HPC” (from “Hahn-Proteinase K-Column”) and “SH” methods. More in detail, while a pH of 11 in the buffer and digestion temperatures of 65 °C for 24 h followed by extraction using the PowerSoil Pro (QIAGEN) kit corresponded to the SH method, a pH of 8 was applied to the HPC one (“Hahn-Proteinase K-Column”) adapted from Hahn et al. (2022) where the authors tailored a commercial column-based DNA extraction method (QIAquick; QIAGEN). Moreover, in the latter (HPC), we increased the digestion temperature from the original 24 h at 55 °C to 24 h at 65 °C as in the SH method. Apart from this, a DNA repair step, originally designed for FFPE samples (NEBNext FFPE DNA Repair v2 Module), was added in run#1 for every DNA extract except for the positive controls. While this repair step was already included in the original SH method (Shiozaki et al., 2021); this did not apply to the other two approaches (Hahn et al., 2022, 2024a and Campos and Gilbert, 2012). Based on our run #1 results, we removed this repair step in samples extracted for run #2 (2017 and 2022 ones). For comparison, in both sequencing efforts (runs #1 and 2), we included both repaired and non-repaired (“original”) DNA extracts for a subset of samples (“repaired vs. original” sample pairs). In this sense, we chose one sample from each sampling year, i.e. those with a high DNA yield (ng□µl□¹ DNA) as a considerably higher amount of DNA is needed for repair (“REP” and “ORIG” samples in Table 3). Finally, the QUIAGEŃs QIAamp DNA mini kit® (“QIA” from now on) was slightly adapted to extract DNA from the positive controls (C+1 and C+2 in Figure 1).

**Table 3.**
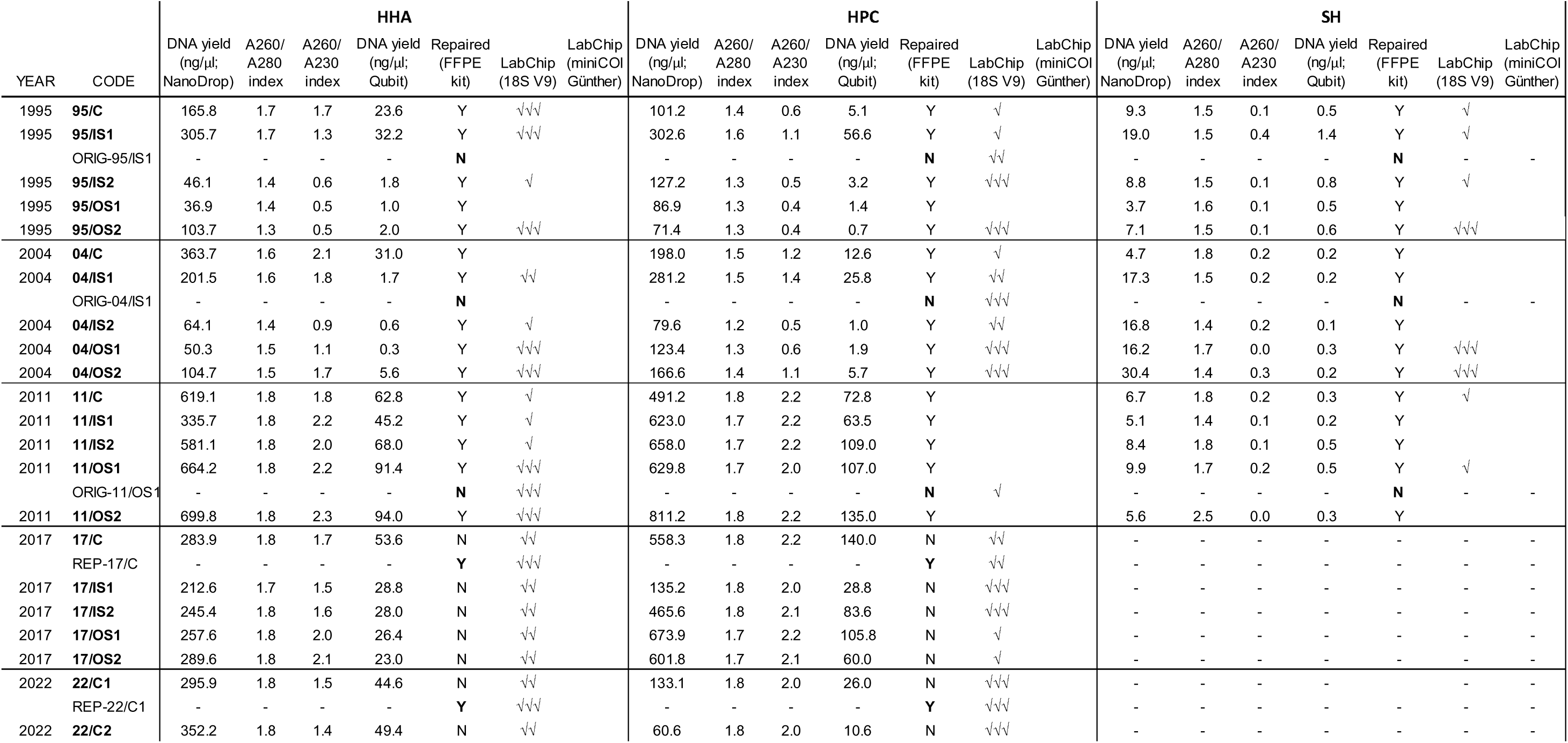
DNA extraction details. DNA yield (both NanoDrop and Qubit measurements), DNA purity indexes (A260/280 and A260/230) and the presence of a band of the expected size in the LabChip are shown for the 22 formaldehyde-fixed samples and the three extraction methods (HHA, HPC and SH), including the repaired (NextNEB FFPE repair kit) / non-repaired (original) sample pairs. Results for both the 18S V9 and the Günther miniCOI marker are shown. See Methods for further details. Fluorescence thresholds for the LabChip band presence assessment: 10-50 units, 50-100 units and > 100 fluorescence units for one, two and three ticks, respectively. Sampleś CODEs as in Table 1.

After extraction, DNA yields (ng□µl□¹) were obtained using two distinct platforms: a NanoDrop ND-8000 spectrophotometer (Thermo Fisher Scientific, Waltham, MA), to measure the concentration of nucleic acids, and a Qubit 2.0 fluorometer (Thermo Fisher Scientific), to measure the concentration of double-stranded DNA. Purity indexes [determined from the absorbance (A) at different wavelengths (nm); A260/A280 and A260/A230 ratios] were also determined with the NanoDrop. Apart from this, for run #1 samples we analyzed DNA integrity (DNA quantification per fragment size; a proxy of fragmentation) by microfluidic capillary electrophoresis in a LabChip® GX Touch HT equipment (High-Throughput Bioanalyzer, Perkin Elmer) utilizing the DNA High Sensitivity Reagent Kit (Perkin Elmer) according to the manufacturer’s guidelines.

DNA repair was performed using the New England BioLabs’ NEBNext FFPE DNA Repair v2 Module following the manufacturer’s guidelines. According to the manufacturer, this kit repairs four types of damaged associated with FFPE samples (deamination of cytosine to uracil, nicks and gaps, oxidized bases and blocked 3’ ends).

Amplicon amplification was tested for a nuclear DNA (nDNA) marker, the 18S rRNA V9 (“18S V9” from now on), and several COI regions of the mitochondrial DNA (mtDNA). Amplicon libraries were constructed following a two-step PCR approach with dual unique indexes (DUIs) of 10 bp (Bohmann et al., 2022). The first PCR added Illumina specific adapters’ overhangs [Forward overhang: 5’ TCGTCGGCAGCGTCAGATGTGTATAAGAGACAG□+forward locus specific primer sequence; Reverse overhang: 5’ GTCTCGTGGGCTCGGAGATGTGTATAAGAGACAG□+ reverse locus specific primer sequence). The second PCR added the required adapters for Illumina sequencing and DUIs. Three different Taq DNA polymerases, namely KAPA HIFI (Roche), GoTaq Flexi (Promega) and QIAGEN Multiplex PCR kit (QIAGEN) were tested; the latter being selected due to its superior performance among tested markers.

Briefly, the 18S V9 region was amplified with the Earth Microbiome primers 1391F-5’-GTACACACCGCCCGTC and EukBr-5’-TGATCCTTCTGCAGGTTCACCTAC (https://earthmicrobiome.org/protocols-and-standards/18s/). This marker, with an amplicon length of ∼150 bp, has shown a remarkable universality among eukaryotes, including the marine plankton assemblage as a whole (de Vargas et al., 2015 and 2022) and also the mesozooplankton community of the BoB (e.g. Albaina et al., 2016; Abad et al., 2016, 2017).

For the COI, the marker of choice for Metazoan barcoding worldwide, due to a combination of high taxonomic resolution and the deepest databases, and also for the zooplankton communities (e.g. Bucklin et al., 2021, Albaina et al., 2024), three regions (namely ‘Leray-Gelle’, ‘Meusnie’ and ‘Günthe’ “miniCOI” markers, with around 313, 130 and 125 bp amplicons, respectively) were tested. However, motivated by our electrophoresis and positive control results, only the latter was finally used for DNA metabarcoding of FF samples. While the Leray-Geller is the most used COI marker with zooplankton community samples (Albaina et al., 2024), Meusnier and Günther ones represents shorter alternatives with, potentially, comparable taxonomic resolution with zooplankton (Zhang et al., 2018 and Günther et al., 2018, respectively):

- The Leray-Geller miniCOI marker was amplified with the primers mlCOIinF-5’-GGWACWGGWTGAACWGTWTAYCCYCC (Leray et al., 2013)/jgHCO2198-5’- GGRGGRTASACSGTTCASCCSGTSCC (Geller et al., 2013). As none of the FF sample tested for run #1 showed the expected size band after the first PCR (apart from the positive control), we discarded this marker from further steps.
- The Meusnier miniCOI marker was amplified with the primers Uni-Minibar F1’-5’-TCCACTAATCACAARGATATTGGTAC/ Uni-Minibar R1-5’- GAAAATCATAATGAAGGCATGAGC (Meusnier et al., 2008). Although originally this marker was coined as universal for eukaryotes, several authors have already warned against its “universal” nature (e.g. Leray et al., 2013). Because of this, we decided to first test marke’s performance with the positive controls. As these failed to amplify most of the copepod taxa, comprising the bulk of the zooplankton in our samples, we discarded this marker from further steps.
- Finally, the Günther miniCOI marker was amplified with the primers nsCOIFo-5’-THATRATNGGNGGNTTYGGNAAHTG and mlCOIintK-5’- GGRGGRTAWACWGTTCAWCCWGTWCC (Günther et al., 2018). This marker was originally designed and validated for North-East Atlantic marine metazoan detection.

For all amplification reactions, 12.5 µl of the QIAGEN Multiplex PCR Kit (QIAGEN), 1 µl of each primer (10 µM), and a variable volume of DNA extract (1–5 µl) were used, for a total reaction volume of 25 µl. Thermocycler programs and additional details are provided in the Supplementary Materials. After an initial electrophoresis (LabChip® GX Touch HT) to evaluate the yield of the first PCR, purification was performed using magnetic beads (CleaNNA NGS kit), followed by the indexing reaction in the second PCR with the Illumina 10 bp unique dual indexes (Illumina® DNA/RNA UD Indexes), following the manufacturer’s standard protocol. Indexed products were assessed by a second electrophoresis, after which amplicons were bead□purified and pooled. Only samples displaying a band of the expected size in the electrophoresis gel (i.e. samples that succeeded amplifying the expected amplicon) were selected for sequencing. However, for the sake of a more comprehensive analysis, in run #1 we opted to (1) include all 18S V9 libraries and (2) include those Günther miniCOI libraries that showed even a faint detectable trace (Table 3, Supp. Mat. Table 1). Sequencing was carried out on an Illumina MiSeq platform using a 150□bp paired□end protocol (MiSeq v2 chemistry). To enhance sequencing performance, library diversity was increased by adding 10% PhiX to the library pool.

### Bioinformatics

The bioinformatic filtering is summarized in Figure 2. Sequence data processing was carried out as described previously by Garate et al. (2022) with minor modifications to compensate for DNA modifications and PCR errors. In a first step, read pairs were overlapped, allowing for up to 40 mismatches, using vsearch v2.14 (Rognes et al. 2016). Primers were then removed using cutadapt v2.8 (Martin 2011) and any overlapped read sequences not containing the full forward and reverse primer sequences, discarded, albeit allowing up to five mismatches. Finally, any merged and primer-free sequences with FASTQ scores suggesting one or more probable errors or lengths falling outside the expected amplicon lengths, once the primer regions are removed, were discarded (100-130 bp for Günther COI, 125-135 bp for Meusnier COI and 125-165 bp for 18S V9 rRNA) using vsearch. Regarding the latter, while amplicon sizes of ∼127 bp were typically observed for Copepoda, some taxa substantially exceeded this length; for instance, Chaetognatha and, in particular, Cladocera, with amplicon lengths of ∼143 and ∼159 bp, respectively. This relatively broad variability in amplicon size among common zooplankton taxa provided an indirect opportunity to infer the influence of DNA fragmentation on the results obtained for this marker. Remaining sequences were de-replicated and sorted by abundance using vsearch and clustered into OTUs using SWARM v3.0 (Mahé et al. 2015) using default settings (-d = 1, fastidious clustering and breaking initial operational taxonomic units likely to represent different sequences). OTU abundances were retained and used to construct an OTU contingency table, discarding singleton OTUs and putative chimeras. Chimeras were predicted using vsearch, reference based with Midori-DARN v253 and SilvaModPR2 v138 as reference database (db) for COI and 18S V9, respectively (https://github.com/lanzén/CREST), followed by *de novo* mode. SilvaModPR2 v138 is a db based on SILVA nr SSU Ref v138 and PR2 v4_13, accessible at https://services.cbu.uib.no/supplementary/crest/. Curated OTUs were aligned to Midori-DARN v253 or SilvaModPR2 v138 using blastn v2.6.0+ and then taxonomically classified using CREST v3.1.0 (Lanzén et al. 2012). Due to its less conserved nature compared to 18S V9 one, we then applied LULU post-clustering curation (Frøslev et al. 2017,) to both Meusnier and Günther miniCOI datasets, with default parameters except for increasing minimum similarity to 97%□, to correct remaining PCR and sequencing artefacts and to merge intra-specific or intra-genomic OTUs.

**Figure 2.**
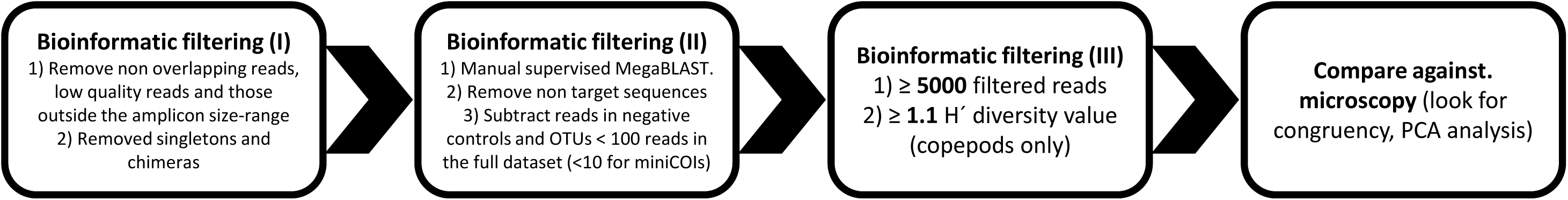
Bioinformatic filtering. Three filtering steps were applied to the DNA metabarcoding results of the formaldehyde-fixed samples (see Methods for further details). Only those samples fulfilling the bioinformatic filters were compared against microscopy. Further details about the number of sequences after each filtering step in Supp. Mat. Table 1.

Moreover, a supervised manual taxonomic assignment step performed by a trained zooplankton taxonomist was included to check the taxonomic assignments of the most abundant OTUs. We focused on those OTUs previously not assigned to species, and aligned them against NCBI GenBank non-redundant nucleotide collection (nr/nt) database with MegaBLAST. More into detail, we manually inspected every OTU assigned above phylum level if it represented > 100 reads; for OTUs below Phylum level, we checked those OTUs with > 5000 reads (but > 1000 for Copepoda). This is critical for 18S V9 where amplicon synonymy (100% identity) has been reported even at the Family level for certain copepod species (whilst, at the same time, being able to discriminate congeneric copepod species; e.g. Albaina et al. 2016). In such cases, providing a 100% amplicon length cover and ≥ 99% identity, species level assignments with MegaBLAST were considered if the biogeography of the assigned species matched the herein sampling area (based on previously published works and/or WoRMS https://www.marinespecies.org/index.php and https://copepodes.obs-banyuls.fr/en/ databases). The aforementioned methodology has shown to improve classification with the relatively short and less variable 18S V9 (e.g. Albaina et al. 2016, Simões et al. 2024), given that in some OTUs, there were several potential classifications over the 99% identity threshold. Moreover, due to SILVA db requirements, short barcodes (< 300 bp) such as those encompassing solely the V9 region are not incorporated (Pruesse et al., 2007), therefore, being NCBI GenBank db their sole repository.

Following taxonomic assignment, we removed non-target sequences (leaving only marine metazoans except for fishes). In this sense, fisheś sequences were removed because most fish eggs and larvae and sorted out just after sampling to compute the DEPM. Besides, as fish specimens are also captured and dissected during the campaign, exogenous fish DNA could be a source of contamination of the plankton samples. Moreover, to minimize contamination issue, we followed Drake et al. (2022) and 1) subtracted the sequences in negative controls from the corresponding OTUs across all the samples, and 2) discarded every OTUs with < 100 reads in the full dataset (this threshold lowered to < 10 for miniCOIs due to its lower availability in the dataset). Finally, only samples with > 5000 reads left and with a copepod Shannon-Wiener diversity (H’) > 1.1 were considered for further analyses (i.e. samples that fulfilled the bioinformatic filtering *sensu* Table 2). The latter is a conservative value based on the H’ values obtained for the biomasses of copepods in the full set of samples as identified under the microscope (Supp. Mat. Figure 1), but also fits with the diversity range reported for marine plankton metabarcoding at a global scale (De Vargas et al., 2022). We used the diversity of copepod biomasses as 1) it is well known that amplicon reads are more comparable to zooplankton biomasses than abundances (e.g. Ershova et al., 2021, 2023), and 2) we could only estimate biomasses for copepods (that represented the bulk of the zooplankton community in the present work: 86% of the counts in the microscopy analyses).

### Statistics

Multidimensional analyses of the sampled stations and the zooplankton taxa were carried out using Principal Components Analysis (PCA) applied to arcsine transformed relative abundance (0-1) values. By comparing microscopy and DNA metabarcoding-based PCA analyses, the congruence of both approached can be assessed (i.e. samples that showed congruency with the microscopy assessment), allowing to identify misperformances (e.g. Blanco-Bercial et al., 2026). The PCA tests were performed using version 5 of CANOCO (Šmilauer and Lepš, 2014). When comparing DNA metabarcoding against microscopy, only those samples fulfilling the bioinformatic thresholds (Figure 2) were included in the PCA analysis; moreover, based in the repaired vs. original sampleś pair analysis, only repaired samples where included from run #1 but original ones from run #2.

## RESULTS

### Microscopy characterization

A total of 58 taxa were identified under the microscopy when considering the 22 FF samples and the two positive controls; 34 of them corresponding to the Class Copepoda (Table 2). Zooplankton abundances in the studied dataset ranged from 2800 to 14800 ind. m^-3^. Copepods represented 85.5% of the abundances for the whole dataset, showing a maximum of 98.7 % (Figure 3a). The minimum values corresponded to the four samples collected in 2022, two coastal stations where two consecutive samples were preserved in formaldehyde (C1 and C2) and in ethanol (C+1 and C+2), with values ranging between 67.4 and 75.1 %. Apart from Copepoda, only other five categories encompassed > 1 % of zooplankton total abundance: Cnidarians (both Siphonophora and other cnidarian taxa; 1.2 and 1.8 %, respectively), Echinodermata larva (2.2 %), Cladocera (1.7 %) and Chordata (4.2 %; all of them appendiculareans). These last two groups comprised five and three distinct taxa, respectively (Table 2). Figure 3b shows the abundance repartition among non-copepod taxa in the sampled stations. After copepods, the most abundant taxa in the 2022 samples were appendiculareans (*Fritillaria* and *Oikopleura* spp.) and echinoderm larvae. Regarding Copepoda, *Oithona helgolandica*, *Oncaea* spp. and the P-Cal category were the most abundant ones with 28, 21 and 19 % of the counts, respectively, being numerically dominant along the whole studied period (Figure 3c). As expected, coastal species (i.e. *Temora longicornis*, *Oncaea* spp., *Euterpina acutifrons, Acartia clausi*) abundances decreased towards the outer stations, the opposite applying to more shelf-oceanic taxa (i.e. Calanidae, *Oithona plumifera*, *Clausocalanus* spp., *Euchaeta* spp.). Regarding copepod biomasses, the highest contributions corresponded to *Calanus helgolandicus* (copepodite stages IV–VI), P□Cal, and *T. longicornis*, accounting for 19 %, 15 %, and 13 %, respectively (Figure 3d). *C. helgolandicus* is the most abundant large copepod (>2 mm in length) in the study area. While this taxon was present along the entire transect in 2004, it was restricted to the outer□shelf stations in 1995 and 2011 and was not detected in 2017, nor in the 2022 samples, which were limited to the coastal zone. Notably, the abundance of *C. helgolandicus* was much higher in 2004, reaching 35–73 % of the copepod biomass across the five transect stations, with the highest values observed at the outer□shelf stations (Figure 3d). This was reflected in the PCA results (Figure 4), where the 2004 stations were displaced relative to their counterparts from other years. This displacement applied only to the outer□shelf stations (OS1, OS2) when using zooplankton abundances, but extended to all five stations (C, IS1, IS2, OS1, OS2) when considering copepod biomass alone. More specifically, in 1995, 2011, 2017, and 2022, stations plotted according to their position along the shelf (from coastal to outer shelf, roughly left to right in Figure 4), regardless of sampling year. In contrast, the 2004 stations—particularly in the copepod□biomass PCA—clustered according to their shelf position but were also clearly separated from stations sampled in the other years. The first two PCA axes explained 48.4% and 61.3% of the variance for zooplankton abundances (Figure 4a) and copepod biomasses (Figure 4b), respectively.

**Figure 3.**
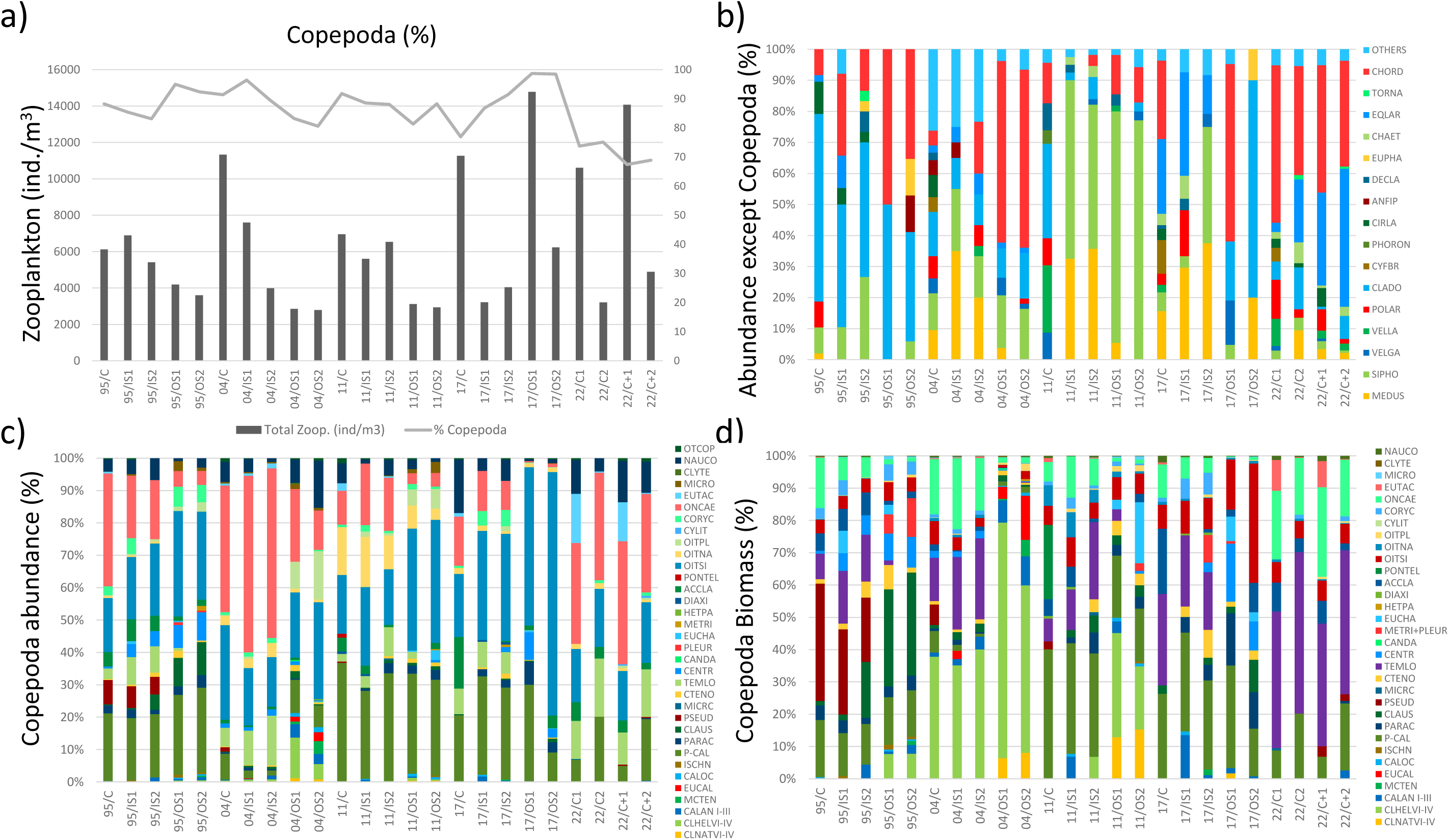
Taxonomic composition (microscopy). Taxonomic characterization of the 24 samples of zooplankton selected for the experiment: in a) absolute zooplankton abundances (ind. m^−3^; black bars, left axis) are shown along with the percentage of copepods (grey line, right axis); relative abundances (%) are shown for b) zooplankton abundance when copepods are removed, c) copepod abundance and d) copepod biomass (mg C dry weight). Sampleś CODEs as in Table 1; taxás CODEs as in Table 2.

**Figure 4.**
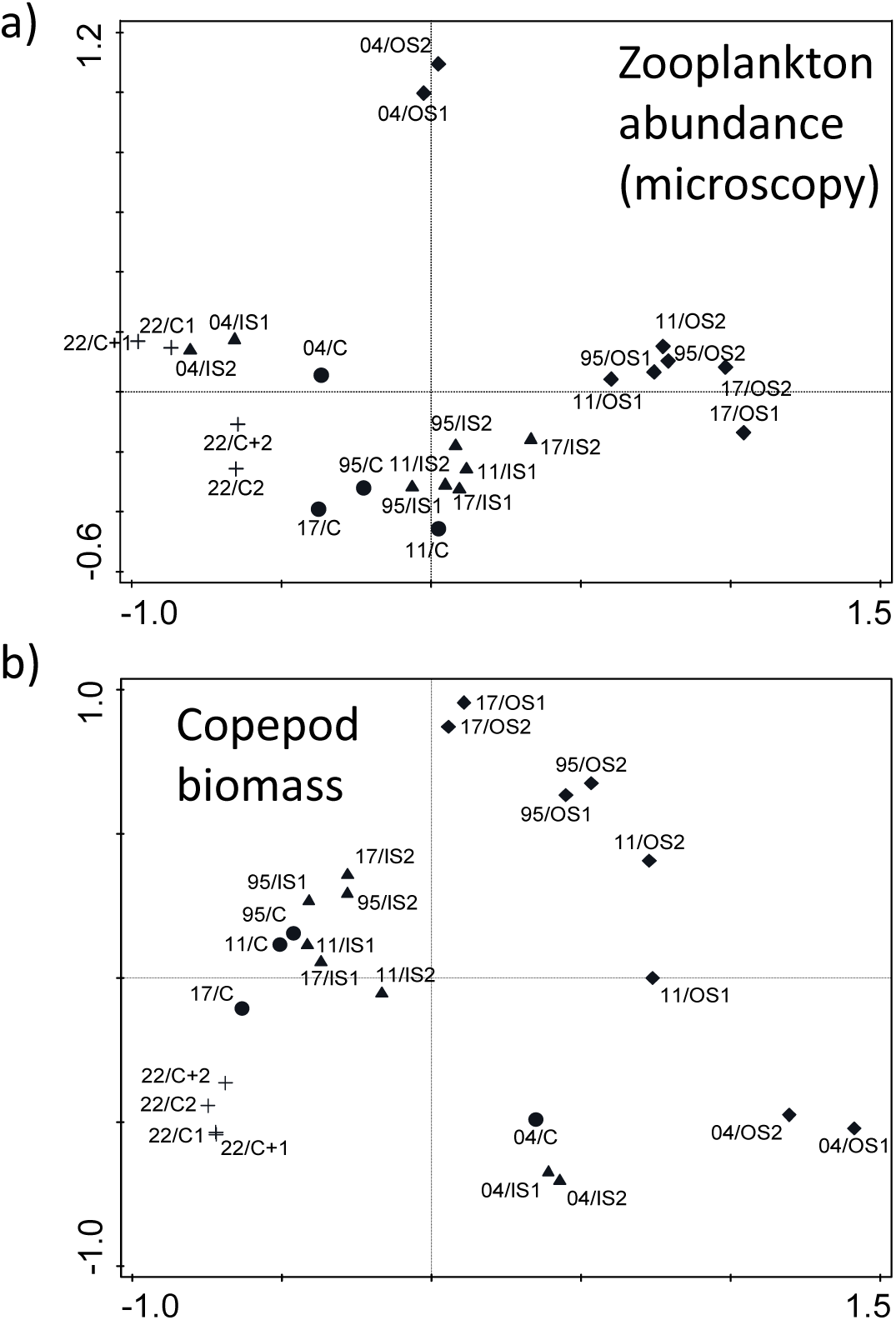
Zooplankton assemblages (microscopy). Principal Component Analysis using arcsine transformed relative abundance (0-1) values of the 24 zooplankton samples selected for the experiment: a) with all the zooplankton taxa and b) based in copepod biomass data only (see Methods for further details). Stationś symbols and labels (CODEs) as in Figure 1 and Table 2, respectively.

### DNA extraction and amplification success of formaldehyde-fixed (FF) samples

Two clear patterns arose for the DNA yield (Table 3): on the one side, the SH method failed to extract DNA (Qubit average values of 0.4 ng□µl□¹ for the 1995-2011 samples with a maximum of 1.4 ng□µl□¹), this confirmed by the DNA integrity results (Supp. Mat. Figure 2), thus discarding this method for the second sequencing effort; on the other side, HHA and HPC methods showed a similar performance (average values of 30.7 and 40.1 ng□µl□¹ for the same period), with DNA yields consistently > 20 ng□µl□¹ from 2011 onwards (except for the 22/C2 sample in the HPC method). These patterns were also followed by the DNA purity indexes, with values well below the ideal range (∼1.8 and between 2-2.2 for A260/280 and A260/230 indexes, respectively) for the SH method, and better purity values, consistently over the ideal threshold from 2011 onwards, for both the HHA and HPC methods.

For the HHA and HPC extracts, as expected, DNA integrity inversely correlated with storage time, with only few samples in 1995 and 2004 showing fragments over the 150-200 bp size range but consistently well over this size in 2011 (Figure 5 and Supp. Mat. Figure 2). Apart from this, a higher fragmentation in the samples extracted with the HHA method when compared with the HPC one was evident for 2011 samples. This corresponds well with the pH values decreasing from the neutral values in the most recent samples (2022 ones; 2 y in formaldehyde) to increasingly acid ones in the rest of the series (Table 1) and with the hasher conditions of the HHA method compared to the HPC one.

**Figure 5.**
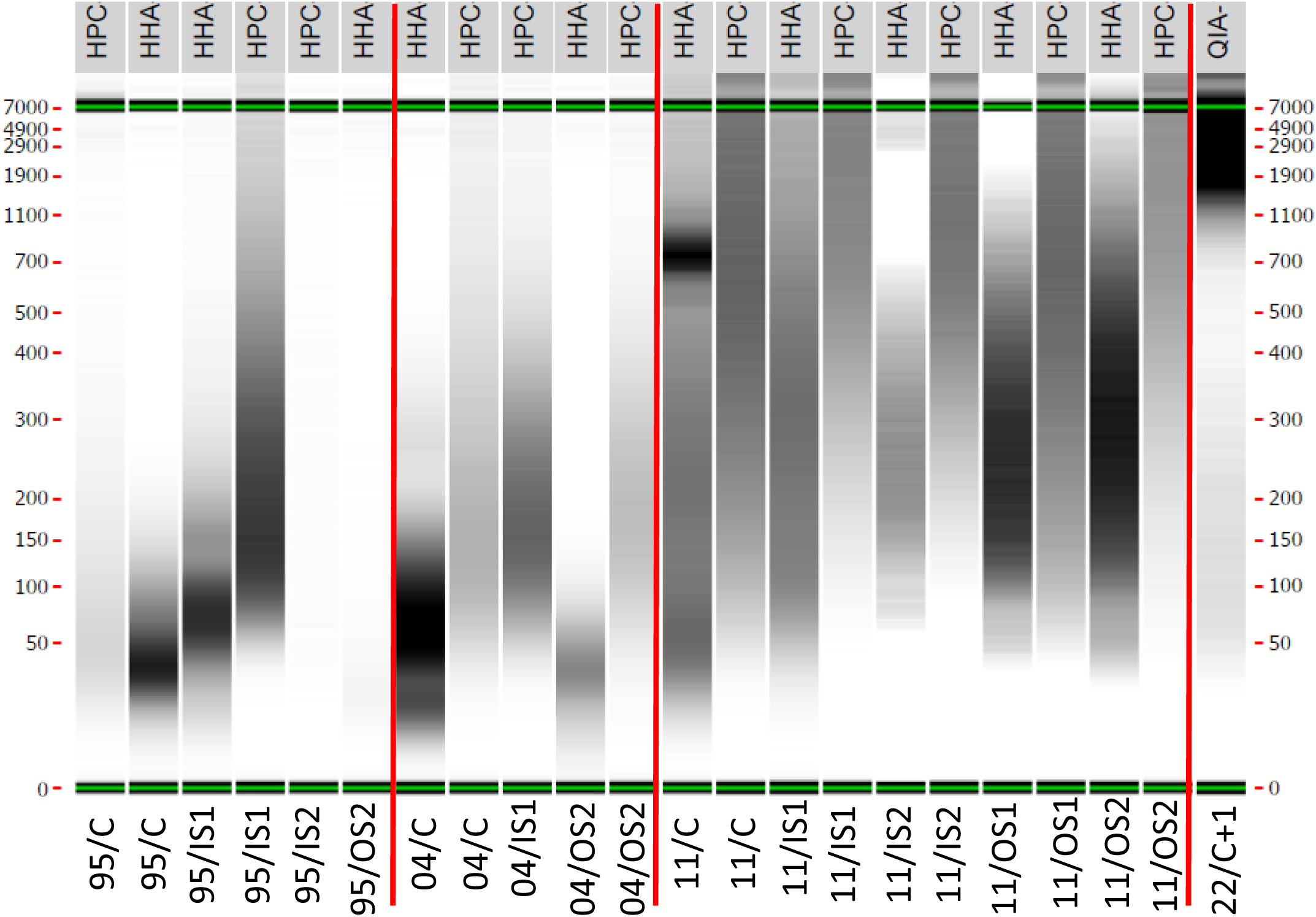
DNA integrity (LabChip). LabChip gel showing the degree of fragmentation for the DNA extracted from 1995-2011 formaldehyde-fixed samples. Only those samples with the higher DNA yields, according to Table 3, are shown here. The rest of samples are in Supp. Mat. Figure 2. Sampleś CODEs in the bottom as in Table 1, extraction methods at the top of the image. A positive control (22/C+1) is shown for comparison.

Regarding amplicon amplification success, a distinguishable band of the expected size was detected only for the 18S V9 marker (Table 3). More in detail, eight out of 15 samples yielded the expected gel band for the SH method, this increasing to 23 and 21 out of 27 (including repaired and non-repaired/original sample pairs) for HHA and HPC methods, respectively. The main difference between HHA and HPC regarding PCR amplification success was the general failure with 2011 samples for the latter; this showing that a high DNA yield does not necessarily correspond to PCR amplification success with FF samples. It also shows that the DNA integrity can affect differently the HHA and HPC amplification success. While, as expected, the fixing time represents a general amplification success predictor for FF samples, DNA integrity would be a good proxy for amplicon amplification success of up to 28 y old samples for the HHA extraction method, but not for the HPC one. Apart from this, the non□repaired (original) pairs of samples from 1995 and 2004 failed to amplify using the HHA method, whereas amplification was successful when using the repaired extracts, as well as for both DNA extract types (repaired and non□repaired) when processed with the HPC method. For the remaining repaired/non□repaired pairs (2011, 2017 and 2022), both methods successfully amplified the expected amplicon, with the sole exception of the repaired extract from 2011 with the HPC method. These results suggest a similar or even superior performance of the non-repaired (original) extracts with samples < 12 y in formaldehyde but the contrary in older samples (at least with the HHA method). Finally, as mentioned in the Methods, the lack of bands of the expected size for the Leray-Geller miniCOI marker in the 1995-2011 samples made us discard this marker.

### DNA metabarcoding of positive controls (ethanol fixed; 18S V9, Günther miniCOI and Meusnier miniCOI markers)

The comparison of the zooplankton assemblages obtained by DNA metabarcoding and microscopic inspection made us discard the Meusnie’s marker for FF samples analysis. More in detail, while copepods dominated the zooplankton community in the positive controls (22/C+1 and 22/C+2 samples; Figure 1) assessed by metabarcoding with both the 18S V9 and the miniCOI Günther markers, as they did when assessed under the microscopy (Figure 3a), these were practically absent from the Meusnier miniCOI one (Figure 6 and Supp. Mat. Figure 3). This confirmed the lack of amplification of the Class Copepoda with the latter, discarding its use with FF samples. Apart from copepods, the most abundant taxa in the positive controls were appendiculareans and echinoderm larvae for the 18S V9 but only the latter for the miniCOI Günther one (Figure 6 and Supp. Mat. Figure 3). The failure of the Günthe’s marker to amplify appendiculareans is confirmed by the microscopy results (Figure 3b). On the contrary, although *Temora longicornis* represented the most abundant copepod for both markers, corresponding with the higher biomass measured for this taxon under the microscope (Figure 3d), the Günthe’s marker showed a superior taxonomic resolution within the Copepoda community (Supp. Mat. Figure 3).

**Figure 6.**
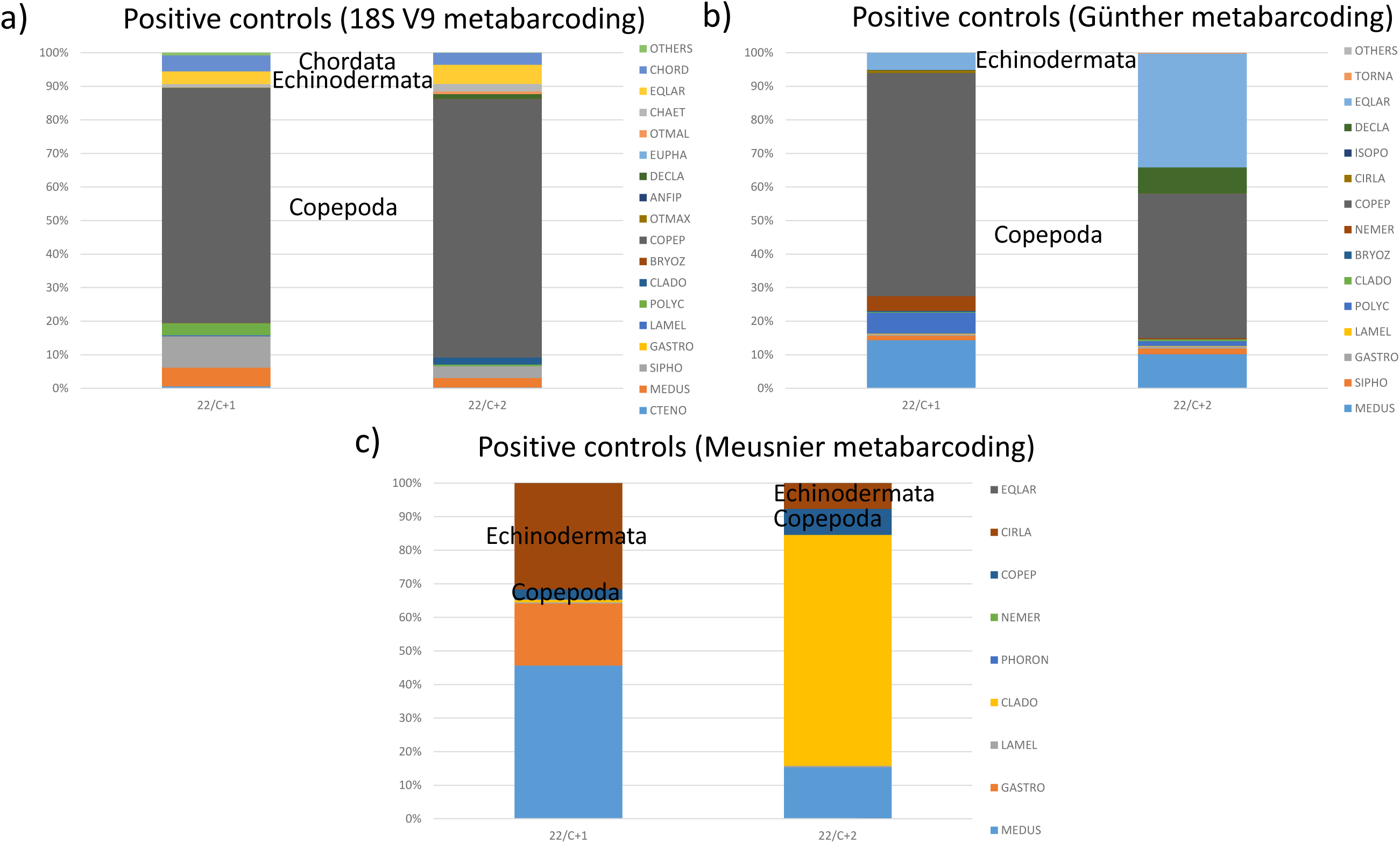
Taxonomic composition of the positive controls (DNA metabarcoding). Taxonomic characterization of the two positive controls (zooplankton samples fixed in ethanol, 22/C+1 and 22/C+2; Table 1 and Figure 1) for: a) 18S V9 b) Günther miniCOI and c) Meusnier miniCOI. Sampleś CODEs as in Table 1. Taxás CODEs as in Table 2 except for the inclusion of Nemertea (NEMER) and Isopoda (ISOPO); moreover, the “OTHERS” category for the Günther miniCOI included Porifera, Phoronida, Annelida and Gastrotricha. Supp. Mat. Figure 3 shows the same graphs but at the highest taxonomic level for each marker.

### DNA metabarcoding of FF samples (18S V9 and Günther miniCOI markers)

None of the FF samples tested for Günther miniCOI metabarcoding passed the bioinformatic thresholds (Figure 2 and Supp. Mat. Table 1), as corresponding with the lack of band of the expected size in the LabChip analyses (Table 3). However, a higher success was reported for the 18S V9 marker (Table 4 and Supp. Mat. Table 1). Interestingly, although necessary, the presence of a band of the expected 18S V9 amplicon size did not guarantee a successful metabarcoding (i.e. samples fulfilling Figure 2 filters). While, as expected, no sample fulfilled this for the SH method (0/15; 1995-2011 period), only nine and seven out of 22 samples (1995-2022 period) did for the HHA and HPC methods, respectively (Table 4; without taking into account the effect of the DNA repair kit, see below).

**Table 4.** Samples fulfilling bioinformatic filters. The number of samples passing the bioinformatic filtering (Figure 2 and Methods for further information) are shown per sampling campaign and extraction method. Only data for the 18S V9 are shown as the other markers did not fulfil the required thresholds. Results only for repaired (NextNEB FFPE repair kit) samples and non-repaired (original) ones for run#1 (1995, 2004 and 2011 samples) and run#2 (2017 and 2022 ones), respectively. Further details about the number of sequences after each filtering step in Supp. Mat. Table 1.

| 18S V9 | 1995 | 2004 | 2011 | 2017 | 2022 |
| --- | --- | --- | --- | --- | --- |
| HHA | 1/5 | 0/5 | 5/5 | 1/5 | 2/2 |
| HPC | 1/5 | 1/5 | 0/5 | 3/5 | 2/2 |
| SH | 0/5 | 0/5 | 0/5 | - | - |

While both methods succeeded metabarcoding the most recent FF samples (2022 C1 and C2; 2/2 for both protocols), they performed similarly poorly with the oldest samples (1/10 and 2/10 for the 1995–2004 samples using HHA and HPC, respectively; Table 4). In contrast, their performance diverged notably for the 2011 and 2017 samples. For the 2011 samples, all extracts met the bioinformatic thresholds under the HHA method (5/5), whereas none met the criteria under the HPC protocol, which was already reflected in the very low number of sequencing reads prior to the second filtering step and consistent with the amplification success results (Table 3 and Supp. Mat. Table 1). Conversely, the 2017 samples showed the opposite pattern: one out of five samples passed the thresholds under HHA, compared with three out of five under HPC (Table 4). Additionally, it is noteworthy that non□repaired (original) samples consistently yielded higher numbers of 18S V9 sequences than their repaired counterparts—on average 93% more reads after Filter□II (Figure 2, Supp. Mat. Table 1).

The fact that a majority of samples showing amplification success (LabChip data, Table 3) failed after bioinformatic filtering (Table 4; Supp. Mat. Table 1) was two-fold: either due to (i) a high proportion of non□metazoan taxa, and/or (ii) extreme dominance by one or a few metazoan taxa. Regarding the first issue, among the 34% of 18S V9 sequences removed during Filter□II, the overwhelming majority (73%) corresponded to fungal taxa. Moreover, a single fungal taxon (*Aspergillus* spp.) in one sample (95/OS2) accounted for 75% of those fungal sequences. The extremely low DNA integrity of this sample (Figure 5), combined with the fact that *Aspergillus* represented 99.4%, 99.6%, and 99.7% of reads in the HHA, HPC, and SH extracts, respectively, suggests contamination occurred either during storage (e.g. during ethanol transfer) or prior to extraction at the sequencing facility. This pattern was also observed with the Günther miniCOI marker, for which the same sample yielded 99.3%, 99.1%, and 99% fungal reads under the HHA, HPC, and SH methods, confirming this interpretation. In this sense, although unintended, the 95/OS2 extracts (HHA, HPC, SH) effectively acted as a type of internal positive control (IPC), indicating the absence of inhibitory processes during extraction and sequencing.

The second major cause of bioinformatic filtering failure was the extreme dominance of one or a few metazoan taxa, which affected those FF samples with >5000 18S V9 reads not passing Filter□II. This subset comprised 15 samples—seven from HHA, six from HPC, and two from SH (the latter restricted to 1995–2011; Supp. Mat. Table 1). More specifically, in samples meeting both bioinformatic criteria (i.e. >5000 sequences after Filter□II and copepod H′ > 1.1; n = 24), the most abundant taxon represented an average of 31% of total sequences. By contrast, in samples meeting only the sequence□number threshold (>5000 after Filter□II; n = 15), this value increased to 80%, clearly demonstrating a sequencing□bias effect in FF zooplankton samples. This observation prompted us to incorporate a biodiversity index as an additional filter in the metabarcoding workflow for FF samples (Figure 2; see Methods for further details).

### Effect of FFPE repair kit (repaired vs. original sample pairs; 18S V9 marker)

From the samples selected to test the effect of the DNA repair step (95/IS1, 04/IS1, 11/OS1, 17/C and 22/C1; see Methods), only the non-repaired (original) DNA extracts of the first two samples, and only for the HHA extraction, failed to amplify and were not sequenced (Table 3 and Supp. Mat. Table 1). Following metabarcoding, while the 2004 sample failed to pass the bioinformatic filtering for both extraction methods, the 2011 one did for the HPC one (Supp. Mat. Table 1). This left us with four pairs of repaired and non□repaired samples suitable for comparison: two shared by both extraction methods (the more recent samples, 17/C and 22/C1) and one exclusive to each method (95/IS1 for HPC and 11/OS1 for HHA) (Figure 7). Across these four comparable pairs, no differences in zooplankton community composition were observed between the repaired and original extracts. Additionally, it is noteworthy that Cladocera—with a longer amplicon (see Methods)—were detected only in the 2017 samples processed with the HPC method.

**Figure 7.**
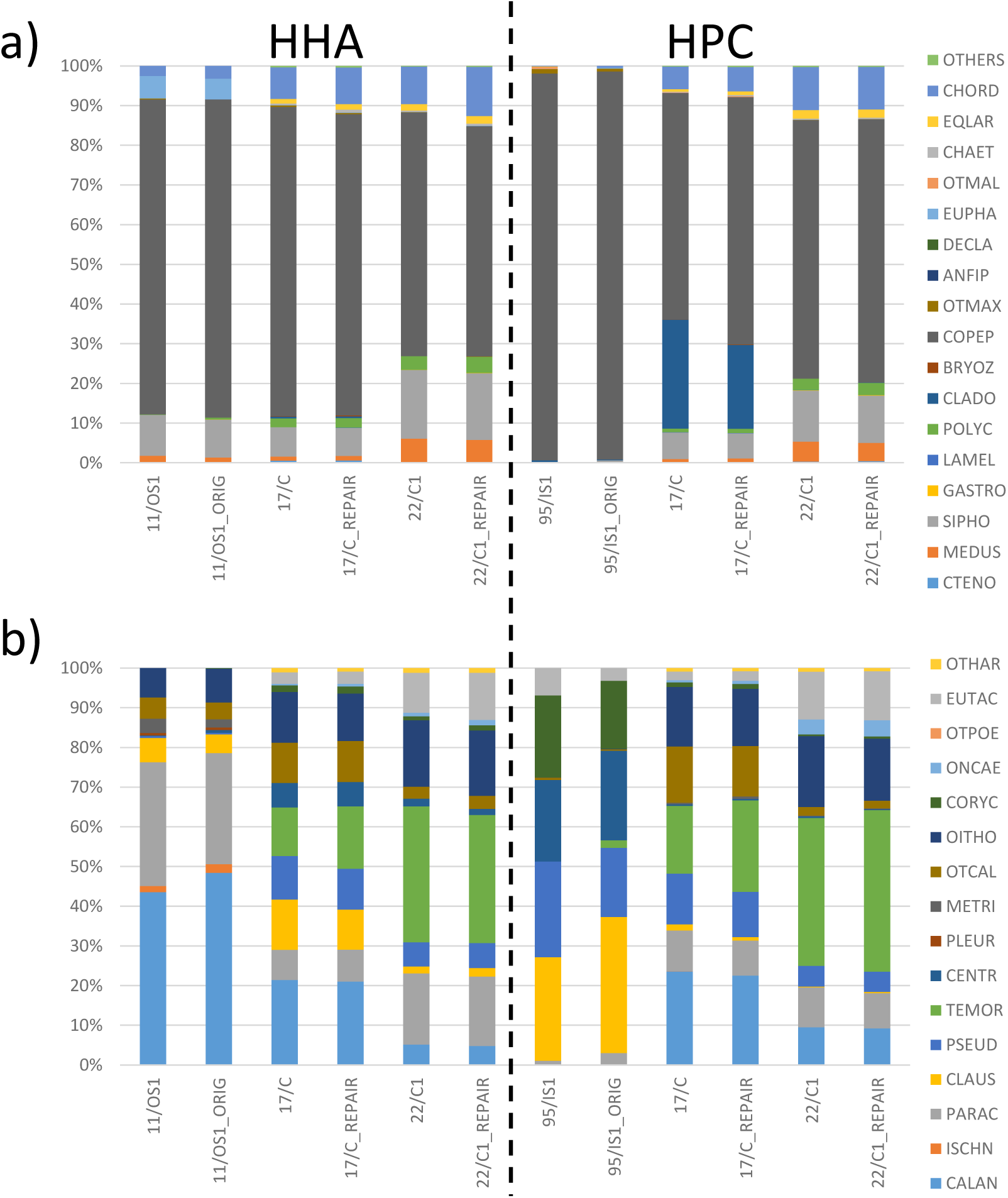
Taxonomic composition of the repaired/non-repaired sample pairs (18S V9 DNA metabarcoding). Taxonomic characterization of the repaired (NextNEB FFPE repair kit) / non-repaired (original) sample pairs for the HHA and HPC methods (left and right side of the graphs, respectively): a) zooplankton 18S V9 sequences (%) and b) copepod 18S V9 sequences (%). Taxás CODEs as in Table 2. Sampleś CODEs as in Table 1; only those samples fulfilling the bioinformatic filters in Figure 2 were included.

### Zooplankton characterization of FF samples via 18S V9 metabarcoding and comparison with microscopy

Due to the absence of differences between repaired and non□repaired extracts, the characterization of zooplankton communities was based on the repaired DNA extracts for run #1 and on the original (non□repaired) extracts for run #2 (see Methods). A total of 37 taxa were identified by 18S V9 metabarcoding in the FF samples, 16 of which belonged to the class Copepoda (Table 2; Figure 8). It should be noted that only four FF samples met the bioinformatic thresholds for both extraction methods (95/IS1, 17/C, 22/C1, and 22/C2). For the remaining samples, metabarcoding results were available for only one extraction method. More specifically, five additional samples passed the thresholds under HHA (all corresponding to the 2011 transect: 11/C, 11/IS1, 11/IS2, 11/OS1, and 11/OS2), whereas three additional samples met the criteria under HPC (04/OS2, 17/OS1, and 17/OS2). In total, 16 FF samples are represented in Figure 8. Across these samples, copepods dominated the 18S V9 sequence composition, followed by Siphonophora and Chordata (appendiculareans), accounting for 68%, 9%, and 8.3% of the reads, respectively (Figure 8a). Among Copepoda, the taxa with the highest proportions of reads were *Temora longicornis*, Calanidae, and *Paracalanus* spp. (20.3%, 19.3%, and 14%, respectively; Figure 8c).

**Figure 8.**
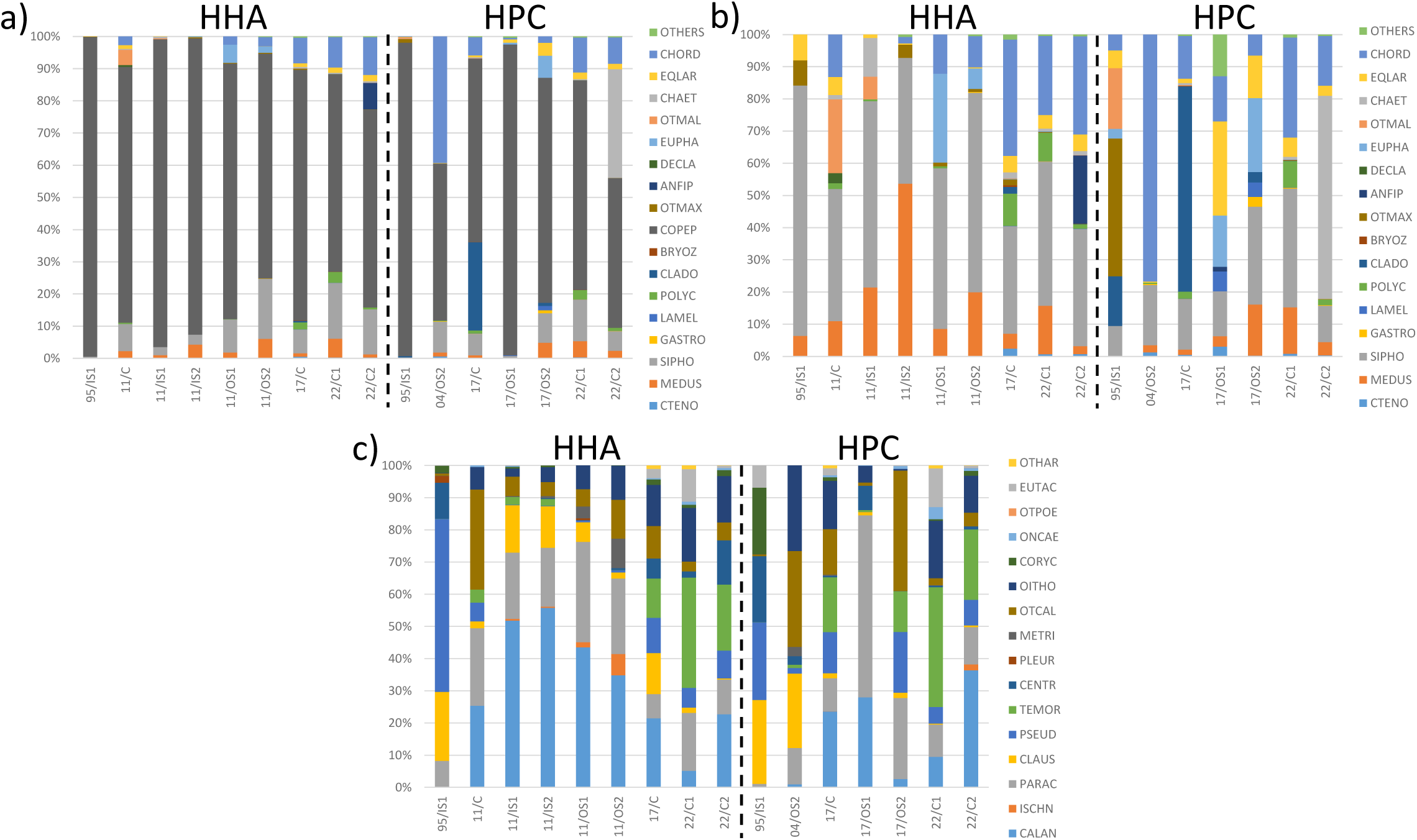
Taxonomic composition of the FF samples (18S V9 DNA metabarcoding). Taxonomic characterization (18S V9 metabarcoding) of the FF zooplankton samples fulfilling the bioinformatic filters in Figure 2. Samples extracted with the HHA and HPC methods are plotted at the left and right side of the graphs, respectively. Relative abundances (%) for a) zooplankton sequences b) zooplankton sequences when copepods are removed and c) copepod sequences only. Only repaired samples for run #1 and original samples for run#2 were included (see Methods for further details). Sampleś CODEs as in Table 1; taxás CODEs as in Table 2.

Figures 9a and 9b show the PCA results for the metabarcoding data of these 16 FF samples, plotted together with the two positive controls (22/C+1 and 22/C+2). The first two PCA axes explained 56% and 50% of the variance for the zooplankton (Figure 9a) and copepod (Figure 9b) 18S V9 communities, respectively. Figure 9c displays the PCA based on microscopy□derived copepod biomass for the same samples (n = 14; the reduction reflecting the four samples for which both extraction methods passed the bioinformatic filters in the DNA metabarcoding dataset). In this case, the first two axes explained 67% of the variance.

**Figure 9.**
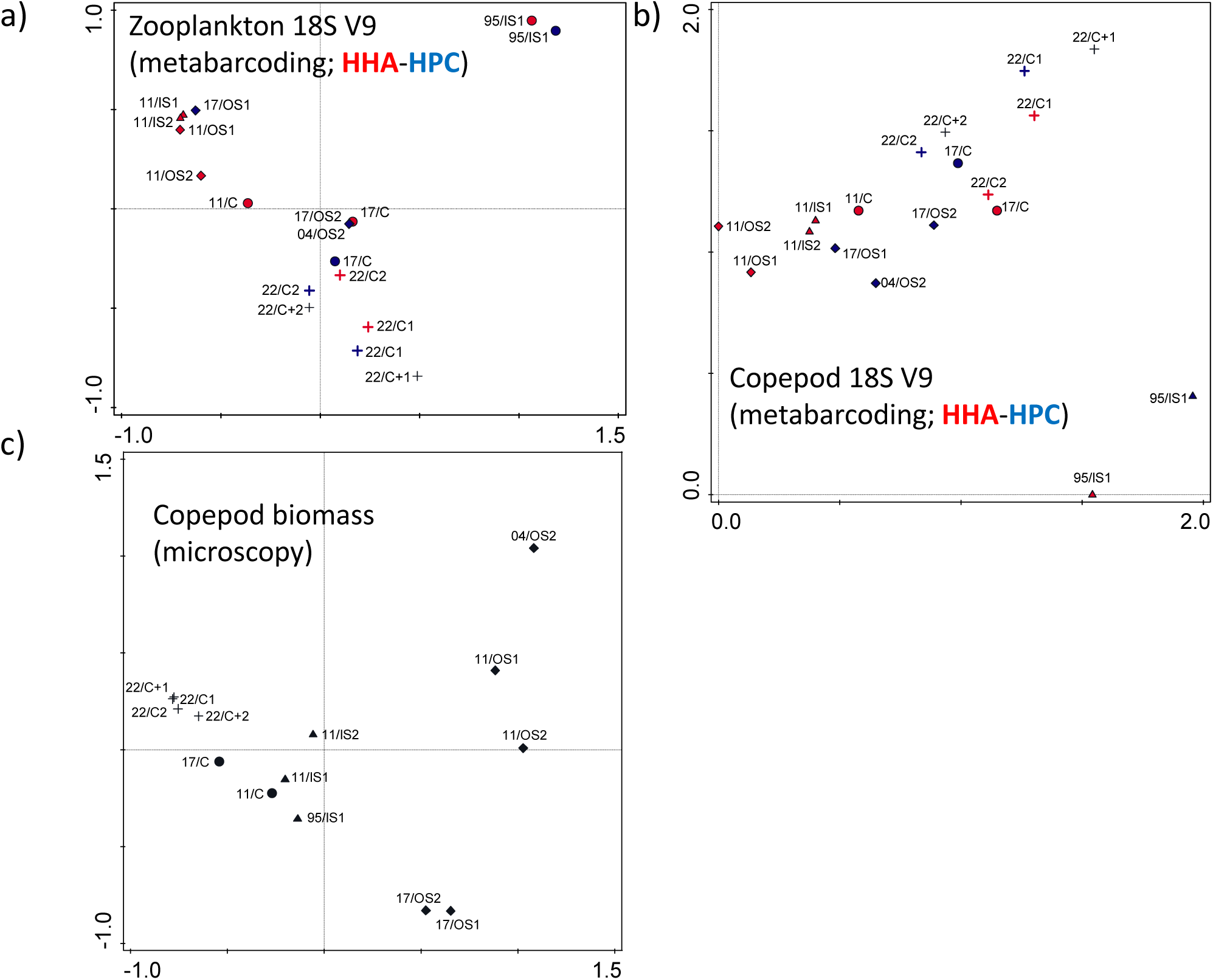
Zooplankton assemblages (18S V9 DNA metabarcoding). Principal Component Analysis using arcsine transformed relative abundance (0-1) values of the zooplankton samples fulfilling the bioinformatic filters in Figure 2. Samples extracted with the HHA and HPC methods are coloured in red and blue, respectively. PCA analyses based on a) zooplankton 18S V9 sequences b) copepod 18S V9 sequences are shown along with c) the PCA for the same samples but based on copepod biomasses (microscopy) for comparison. For the FF samples only repaired samples for run #1 and original samples for run#2 were included (see Methods for further details); positive controls (ethanol fixed samples: 22/C+1 and 22/C+2) were also added to the analysis. Stationś symbols and labels (CODEs) as in Figure 1 and Table 2, respectively.

Whereas the coastal□to□outer□shelf gradient remained evident in the microscopy□based PCA—even for this reduced subset of samples (Figure 9c; see also Figure 4b for the full dataset)—this pattern was largely absent in the metabarcoding□based ordinations for FF samples, both for the zooplankton and the copepod communities (Figures 9a and 9b, respectively). More specifically, among the metabarcoding samples considered, only the most recent ones (22/C1 and 22/C2) showed the expected spatial structure, clustering with their positive□control counterparts (22/C+1 and 22/C+2) under both extraction methods (Figure 9a and b) and aligning well with the microscopy□based patterns (Figures□4,□9c). Beyond these, only the 2017 samples (17/C for both HHA and HPC, and 17/OS1 and 17/OS2 for HPC) showed a subtle correspondence with their position along the sampled transect. In contrast, the samples from 1995, 2004, and 2011 clustered independently of their position along the shelf, indicating a poor performance of metabarcoding relative to microscopy for these older FF samples. In particular, the five samples from the 2011 transect clustered tightly together and distinctly apart from the rest. Similar mismatches were observed for the oldest samples, 95/IS1 and 04/OS2. While both HHA and HPC extracts of 95/IS1 formed a clear outlier when compared with microscopy□based ordination, the 04/OS2 (HPC) sample did not exhibit the displacement observed in the microscopy data (likely reflecting the elevated Calanidae biomass in 2004 described earlier; Figures 3d and 4b).

## DISCUSSION

### “Is DNA metabarcoding an option for formaldehyde-preserved zooplankton time series?”

Based on our results, the answer is “it depends.” On one hand, we demonstrate that short, amplifiable DNA barcodes can be recovered by metabarcoding from samples preserved in formaldehyde at room temperature (RT) for up to 28 years (Table 4). This accords with Ripley et□al. (2008), who sequenced a ∼100□bp amplicon of a coccolithophore from formaldehyde fixed (FF) samples up to 35 years old. On the other hand, robust characterization of zooplankton communities by metabarcoding depends on accurate and unbiased sequencing of the whole assemblage, not merely presence–absence of selected taxa. In this respect, 18S V9 metabarcoding of FF zooplankton performed comparably to microscopy only for samples preserved ≤□2 years, with some but inconsistent congruence for 7 years old samples (Figure 9). Given the low success with older material, further method development is needed to unlock the scientific value of long-term archives. We provide recommendations at the end of the Discussion. Importantly, the scope of our findings should extend to other long term zooplankton series preserved in 4% buffered formaldehyde and stored at RT, including major global programs (e.g. CPR since 1931, CalCOFI since 1949, Plymouth L4 since 1988, and IEO CSIC RADIALES since 1990; see https://www.st.nmfs.noaa.gov/copepod/metabase/).

### Factors determining metabarcoding success with FF samples

As anticipated, fixation time was the best predictor of PCR amplification success (i.e. presence of a LabChip band at the expected size; Table□3). However, amplification alone did not guarantee that FF samples would pass bioinformatic thresholds. Of 22 FF samples, 20 (HPC) and 17 (HHA) yielded the expected band, yet only 9 and 7 passed bioinformatic quality filters, respectively (Tables□3, 4; Supp. Mat. Table□1). The main reasons for failure were: (i) a high proportion of non-metazoan reads, and/or (ii) extreme dominance by one or a few metazoan taxa, which rendered even samples with >100□k reads of limited ecological value (Supp. Mat. Table□1). Moreover, even when FF samples fulfilled bioinformatic thresholds, congruence with microscopy was restricted to ≤□2 year samples (HHA and HPC), and only partially evident for 7 year old samples using HPC (Figure□9).

As expected, the lower pH measured in older samples (Table□1) coincided with reduced amplification success, consistent with the acid catalyzed fragmentation of DNA that persists even when formaldehyde induced DNA–protein cross-links are reversed (e.g. Hahn et□al., 2022; Holleley &□Hahn, 2025). Although absolute pH measurements in formaldehyde can be unreliable (e.g. Simmons, 2014), the progressive acidification with storage time is unambiguous (Table□1). Increased fragmentation is supported by three lines of evidence: (a) failure to amplify the 313□bp Leray–Geller miniCOI, (b) declining DNA integrity with storage (Figure□5; Supp. Mat. Figure□2), and (c) loss of the longer 18S V9 amplicons in the oldest samples (e.g. Cladocera in 95/IS1; Figures□3,□8). Although samples were buffered with borax, buffer capacity declines over long storage (Simmons, 2014). Routine pH checking would be ideal, but is operationally impractical for monitoring programs handling hundreds to thousands of samples annually.

Several studies with FF samples show that elevated temperature and/or alkaline pH during lysis help reverse cross-links at the cost of additional fragmentation (e.g. Gilbert et□al., 2007; Totoiu et□al., 2020). Two general approaches exist:

1. Hot alkaline lysis (HA): brief exposure (≤□1□h) to 90–120□°C in strong alkali (e.g. Zhang et□al., 2010; Campos &□Gilbert, 2012; Hahn et□al., 2022,□2024a). Our HHA method followed Campos &□Gilbert (2012) (pH□12–13; 100□°C for 40□min), instead of 120□°C for 25□min (Hahn et□al., 2022,□2024a).
2. Prolonged proteinase□K digestion: typically 24□h to several days, often at 65□°C (e.g. Shiozawa et□al., 1992; Palero et□al., 2010; Gilbert et□al., 2007; Shiozaki et□al., 2021; Straube et□al., 2021). Our SH method used pH□11 at 65□°C for 24□h; our HPC modified Hahn et□al. (2022) increasing temperature from 55 °C to 65□°C for 24□h (pH□8), followed by a column based workflow enabling automation.

Limited exposure to heat + alkali efficiently breaks cross-links (Campos &□Gilbert, 2012), but entails a trade-off: the HHA improves access for polymerase yet exacerbates fragmentation, potentially preventing barcode recovery (Gilbert et□al., 2007). Conversely, milder protocols (HPC)—lacking strong alkali and heat shock—tend to work better with recent samples, where cross-link density is lower. Our results reflect this: HHA was the only method to amplify 2011 samples, whereas HPC outperformed HHA for samples ≤□7 years (Table□4) and was the only method to recover the longer Cladocera amplicon in 2017 (17/C sample; Figure□8). Moreover, we do not recommend a general DNA repair step. Across paired extracts (repaired versus non-repaired/original), we found no significant differences in community composition (Figure□7), while DNA repair reduced yield due to added purification, especially in long fixed samples (Table 3 and Supp. Mat. Table 1; e.g. Steiert et□al., 2023). Although base modification artefacts are a concern, their impact appears limited here and is debated (see Schander &□Halanych, 2003 for a review). In this sense, Steiert et al. (2023) recently reported that FFPE samples are more prone to mutate than samples in a formaldehyde solution as the ones in zooplankton time series. Notably, two of the oldest samples (95/IS1, 04/IS1) amplified with HHA only when repair was included; thus, including a repair step would be recommended in samples preserved for >□11 years. In summary, the HHA method with a repair step performed relatively better with older FF material (> 7□years), albeit with low congruency with microscopy, whereas HPC, without a repair step, would be simpler, automatable, and preferable for more recent samples (≤□7□years).

Other alternatives, not tested in this study, include the use of semi□nested PCR (e.g. Zhang et al., 2010) and single□stranded PCR (Straube et al., 2021; Hahn et al., 2022). In the former approach, Zhang et al. (2010) applied a 30□minute heat shock at 90□°C in a pH□9 lysis buffer to individual fish specimens preserved in formaldehyde for up to 23 years, achieving high amplification success rates ranging from 72% in one□year□old samples to 31% in 23□year□old specimens. Although these ancient□DNA□oriented methods can improve the recovery of short DNA fragments from FF material, they are generally optimized for individual specimens and are therefore unlikely to be practical for bulk community samples. Besides this, Totoiu et al. (2020) successfully amplified a 180□bp mtDNA fragment from individual crabs fixed in formaldehyde for up to 2 years, using a Vortex Fluidic Device (VFD) during digestion to avoid exposure to extreme temperature or pH conditions and thereby minimize further DNA degradation. However, the authors did not assess specimens with longer fixation times, limiting conclusions about their applicability to older archival material. Finally, alternative cross□link removal strategies that do not rely on high temperatures, prolonged digestion, or alkaline conditions—such as those proposed by Karmakar et al. (2015)—are promising avenues for improving the performance of metabarcoding protocols on FF material. Nonetheless, these methods remain untested in zooplankton community samples, and their suitability for complex bulk assemblages has yet to be demonstrated.

### The SH method (Shiozaki et al., 2021) did not work on FF zooplankton samples

Shiozaki et□al. (2021) reported recovery of a ∼450□bp rRNA (18S V7–8) amplicon from zooplankton preserved in formaldehyde at 4□°C for up to 23 years, with microscopy–metabarcoding congruence for diatoms (zooplankton microscopy was not assessed). However, their protocol (‘SH’; lysis with pK in pH□11 and 24□h at 65□°C) failed on all our samples, yielding very low DNA (Qubit ∼0.4□ng□µl□¹ on average; max 1.4□ng□µl□¹) and no visible bands (Table□3), indicating inefficient extraction under our conditions. The most plausible explanation is storage temperature: our samples were kept at RT, not 4□°C. This could be related to the precipitation of formaldehyde at these temperatures (UNESCO 1976, Simmons 2014). In this regard, low temperature storage reduces formaldehyde associated DNA damage (e.g. Koshiba et□al., 1993; Schander &□Halanych, 2003). However, long-term 4□°C storage is logistically and financially unfeasible for many monitoring programs managing thousands of samples (e.g. BIOMAN surveys: 37 campaigns by 2025; >750 samples per campaign).

### Neither DNA yield nor integrity alone predict metabarcoding success in FF samples

Although amplification success in FF samples does not guarantee a community characterization consistent with microscopy, identifying reliable proxies for PCR success before investing time, financial resources, and precious sample material in library preparation and sequencing is essential. Although DNA yield (ng□µl□¹; Table□3) is a traditional indicator of extraction success, our results (and prior work: Palero et□al., 2010; Straube et□al., 2021) show that yield is a poor predictor of PCR or bioinformatic success in FF samples, especially for older samples. Interestingly, Palero et□al. (2010) proposed DNA integrity as a better proxy; indeed, for HHA, integrity predicted amplification better than yield, whereas for HPC, even apparently adequate integrity did not ensure success (2011 samples failure; Figure□5, Tables□3, 4 and Supp. Mat. Table□1). The contrasting performance of the HHA method with the 2011 samples is to be related with a more efficient removal of protein-DNA cross-links associated with formaldehyde fixation of this hasher extraction method. This would be related with the abovementioned trade-off between yielding a less fragmented DNA or a more “amplifiable” one, without cross-links preventing PCR in FF tissue. In summary, we would recommend not using DNA yield as a proxy for PCR success with FF samples but considering integrity screening for HHA workflows.

### 18S V9 vs. miniCOI markers with FF samples

Although our adapted methods recover DNA from 4% buffered formaldehyde at RT even after 28 years, formaldehyde associated fragmentation is irreversible. Consequently, FF time series require short amplicons, ideally ≤□150–200□bp (e.g. Bucklin &□Allen, 2004; Gilbert et□al., 2007; Baird et□al., 2011; Totoiu et□al., 2020; Rowena Stern, personal communication). Our results support this: while Leray–Geller maker (∼313□bp) failed for 1995–2011, the ∼127□bp 18S V9 amplicon was detected, even in the oldest samples, after bioinformatic filtering (Table□4; Supp. Mat. Table□1). Moreover, we also recovered the longer Cladocera 18S V9 amplicon (∼159□bp) in the most recent samples (2017 via HPC; 2022 via both HHA and HPC), consistent with the HHA-HPC trade-off discussed above.

By contrast, the Günther miniCOI (∼125□bp) failed to amplify in all FF samples (Supp. Mat. Table□1). Although this marker offers higher taxonomic resolution than 18S V9, at least for Copepoda (Supp. Mat. Figure□3), the 18S V9 marker, when complemented with expert□curated taxonomic reassignment (see Methods), provided a level of resolution comparable to microscopy (Table□2). Despite its lower intrinsic taxonomic resolution relative to COI□based barcodes, the broad universality of 18S V9 makes it particularly suitable for analysing high□diversity plankton communities (e.g. de Vargas et□al., 2015, 2022; Abad et□al., 2017; Bucklin et□al., 2019). Furthermore, given the known limitations of miniCOI markers for amplifying tunicates/appendicularians and other key zooplankton groups (see Albaina et□al., 2024 for a review), combining a short miniCOI fragment with the more conserved 18S V9 remains an advisable strategy.

The Meusnier miniCOI (∼130□bp) proved unsuitable for marine invertebrates/copepods in our material (Figure□6; Supp. Mat. Figure□3), likely due to primer design issues previously noted (Leray et□al., 2013; Elbrecht &□Leese, 2017), despite some success in single species barcoding (Zhang et□al., 2018). Our positive controls suggest primer-template mismatches across key copepod taxa; we therefore discourage its use on bulk zooplankton. Regarding Günther miniCOI, total failure in our FF samples could reflect the high primers degeneracy (15 ambiguities) and/or reduced accessibility of mtDNA relative to multi-copy nuclear rDNA in FF tissues. While a greater damage to mtDNA than nDNA is expected due to the lack of histones in the former (e.g. Rong et□al., 2021; Liao et□al., 2022), formaldehyde also crosslinks mtDNA associated proteins (Kaufman et□al., 2000). Notably, Díaz Viloria et□al. (2005) failed to amplify even a 176□bp mtDNA target from FF fish larvae (≥□8□months) using nine extraction protocols (but without high heat/alkali conditions). Whether Günther’s failure stems primarily from mtDNA location versus primer degeneracy remains unresolved. To further explore this issue, we evaluated the amplification success of the Meusnier miniCOI marker, slightly longer than the Günther miniCOI (approximately 130□bp versus 125□bp), using the 2017 and 2022 samples. The LabChip results (Supp. Mat. Figure□4) showed a band of the expected amplicon size (using the fluorescence thresholds described in Table□3) in four out of eighteen samples. Among the fourteen non□repaired extracts, only sample 22/C1 (HPC) yielded a band, whereas three of the four repaired extracts amplified successfully (17/C for both HPC and HHA, and 22/C1 for HHA). This pattern suggests that the Meusnier marker may exhibit higher amplification success than the Günther miniCOI. However, as demonstrated previously, amplification success alone does not guarantee that a sample will pass bioinformatic filtering or produce results congruent with microscopy. Although the amplicon sizes of the two markers are comparable, the substantially higher number of ambiguous bases in the Günther primer set likely contributes to its poorer performance. Nonetheless, as noted earlier, the failure of the Meusnier marker to amplify copepods and other zooplankton taxa (Figure□3; Supp. Mat. Figure□3) ultimately limits its utility for metabarcoding FF zooplankton samples.

In short, 18S V9 is promising for FF samples but should be complemented by a short mtDNA marker. Although mtDNA may be less amplifiable than multi copy nDNA in FF, our preliminary test with the Meusnier marker would indicate that ≤□150□bp mtDNA targets are feasible in recent samples if primers are truly universal and minimally degenerate. For instance, Pérez et□al. (2005) amplified a ∼155□bp 16S fish barcode from formaldehyde fixed eggs/larvae stored at RT for up to 10 years using non degenerate fish specific primers. However, the COI region remains the preferred mitochondrial target due to comprehensive reference databases, even for zooplankton (Andújar et□al., 2018; Bucklin et□al., 2021), but we are not aware of any marker fulfilling those requirements for the COI region (e.g. Elbrecht et al., 2019). We therefore recommend: (1) exploring other mtDNA regions, and (2) using cocktails of group specific primers targeting short regions (e.g. Berry et□al., 2023) to complement 18S V9.

### Alternative storage strategies

In many zooplankton time□series programmes, including the one examined here, samples are stored in formaldehyde (formalin) for their entire archive life to ensure long□term morphological stability and taxonomic consistency. Yet, we are aware that other long□term zooplankton series employ an alternative preservation approach commonly used in museum protocols for individual specimens: an initial fixation in formaldehyde for several days or weeks followed by long□term storage in 70% ethanol (e.g. Suthers et□al., 2019, for zooplankton; Simmons, 2014, for a review). Although transferring samples to ethanol represents standard “best practice” in museum curation, laboratory safety, and genetic work, routine monitoring programmes typically prioritize taxonomic consistency across decades, which is more readily achieved by maintaining samples continuously in formaldehyde. Interestingly, in the present study we found that, using both the HHA and HPC extraction methods, DNA metabarcoding of samples preserved in formaldehyde for two years at RT produced results comparable to microscopy. This suggests that, if transfer to ethanol effectively halts or at least reduce further DNA degradation (e.g. Zimmermann et□al., 2008), the methods tested here could hold promise for recovering archived zooplankton biodiversity from time□series in which FF samples are eventually stored in ethanol. In this regard, Kirby &□Lindley (2005) reported recovery of relatively long DNA amplicons (∼500–600□bp) from recently fixed CPR zooplankton specimens; moreover, laboratory studies have reported high sequencing success after very short formaldehyde exposure (e.g. fish eggs, ∼95% success for a 570□bp mtDNA target: Mozfar et□al., 2024). Apart from this, museum specimens fixed briefly in formaldehyde and stored in ethanol for decades have also yielded 470–570□bp mtDNA amplicons (Shedlock et□al., 1997). In contrast, when fixation times are extended, amplification success declines sharply: Zimmermann et□al. (2008) reported complete amplification failure after specimens had remained in formaldehyde for one year prior to transfer to ethanol. In this regard, transfer to ethanol in zooplankton FF samples typically occurs after several weeks of initial fixation, followed by long□term storage in ethanol.

However, previous studies have reported continued DNA fragmentation in formaldehyde□fixed samples stored long□term in ethanol (e.g. Shiozawa et□al., 1992). This has been attributed to the progressive acidification of the solution during ethanol storage, which accelerates DNA degradation (Simmons, 2014; Straube et□al., 2021). Although 70% ethanol is generally preferred over 96% or absolute ethanol for storing formaldehyde□fixed zooplankton, owing to reduced brittleness of the specimens, it is nonetheless slightly acidic (Suthers et□al., 2019). In line with this, Chakrabarty et□al. (2006) working with museum fish specimens concluded that only mtDNA amplicons <□200□bp, and at low success rates, could be obtained from formaldehyde□fixed fish subsequently stored long□term in 70% ethanol at room temperature. Similarly, Díaz□Viloria et□al. (2005) reported that fixation periods exceeding 48 hours prevented amplification of mtDNA barcodes as short as 176□bp from marine fish larvae stored for eight months in 70% ethanol. In contrast, Baird et□al. (2011) showed that, with insect specimens, 17.5% of samples fixed briefly in formaldehyde and stored in 70% ethanol for up to 23 years yielded the short Meusnier miniCOI fragment. Notably, the authors also suggested that long□term ethanol storage might itself exacerbate DNA damage. More recently, Appleyard et□al. (2021), working with individual fish specimens including larvae, demonstrated a combined effect of formaldehyde fixation duration and subsequent ethanol storage time on the amplification of mtDNA barcodes (200–650□bp). Specifically, for samples stored >□3 years in ethanol, amplification success was restricted to the shortest barcodes, and only when fixation had been <□8 weeks. These findings emphasize that the interplay between formaldehyde fixation duration and ethanol storage time critically influences DNA recoverability for barcoding purposes.

Despite these insights, the implications of a brief formaldehyde fixation followed by long□term ethanol storage remain insufficiently understood for bulk zooplankton community samples, and controlled experiments are still needed. Furthermore, Blom (2021), reviewing studies that lacked a prior formaldehyde□fixation step, strongly cautioned against the use of ethanol for both short□ and long□term storage because of its potential to fragment DNA. To our knowledge, the potential additional damage incurred during ethanol storage after formaldehyde fixation has not yet been rigorously assessed for zooplankton samples.

## Conclusions and recommendations

Long□term zooplankton time series are fundamental for tracking and understanding ecosystem change (e.g. O’Brien et□al., 2017; Ratnarajah et□al., 2023; Grigoratou et□al., 2025; Holland et□al., 2025). Given the high turnover rates and strong environmental sensitivity of zooplankton, timely biological indicators are essential for adaptive ecosystem management (Edwards et□al., 2010). Recent works with FF museum specimens have succeeded assembling whole genomes from very short reads (∼60–100□bp) and have even enabled “historical epigenetics” (Hahn et□al., 2024a,b; Holleley &□Hahn, 2025). However, community samples metabarcoding demands longer amplicons to ensure taxonomic resolution (e.g. Meusnier et□al., 2008; Elbrecht et□al., 2019). To our knowledge, only Shiozaki et□al. (2021) attempted DNA metabarcoding of FF zooplankton samples; our results suggesting their success may hinge on cold storage. Here, we focused on the most common scenario for zooplankton time series: long□term storage in buffered formaldehyde at room temperature. Although the HHA and HPC extraction methods were capable of amplifying DNA from samples preserved for up to 28 years, the irreversible fragmentation associated with prolonged formaldehyde exposure imposes significant constraints on metabarcoding applications. Two practical implications of our work follow: (i) short amplicons (≤□150–200□bp) were essential, and (ii) while the presence of target taxa could still be detected in very old FF zooplankton samples, assemblage□level congruence with microscopy was restricted to samples preserved for ≤□2–7 years.

Nearly two decades ago, Bucklin and Allen (2004) anticipated that the molecular characterization of FF zooplankton could unlock century□old archives for population genetic and systematic analyses. While this goal remains only partially realized, recent advances in molecular techniques have allowed meaningful progress. As the present study represents an initial step toward implementing DNA metabarcoding in FF zooplankton time series, we conclude by providing a set of recommendations aimed at guiding and promoting further research in this area:

1. Extraction method selection by fixation time. Use the HHA method with the DNA repair step for samples preserved in formaldehyde for more than seven years, prioritizing those with sufficient DNA integrity before proceeding to library preparation. For more recent samples (≤□7 years), the HPC method without repair is preferable, as it is simpler, automatable, and performs better under these conditions.
2. Target fixation times for ecological analyses. Focus on samples preserved ≤□7 years when the goal is to infer zooplankton community composition. We recommend validating metabarcoding□based assemblages with parallel microscopic identifications for a subset of samples, especially those older than two years.
3. Marker choice and complementarity. Combine the 18S V9 marker with at least one additional universal marker of higher taxonomic resolution and comparable amplicon size. Although short miniCOI markers would be ideal due to their resolution and extensive reference databases, the miniCOI variants tested here failed with FF samples. Thus, there is a need to design new universal mtDNA markers suitable for FF material. In the interim, cocktails of group□specific primers represent a practical alternative.
4. Improvement of reference databases and taxonomic assignment. Continue expanding 18S V9 reference databases for key zooplankton taxa and incorporate expert□supervised taxonomic reassignment to partially overcome the resolution limitations of this marker.
5. Assessing alternative storage protocols. Evaluate the performance of the adapted HHA and HPC methods on samples transferred to ethanol after several weeks or months in formaldehyde. Although not routine practice, some monitoring programmes already employ this strategy and could serve as test cases. Additionally, further testing of FF samples stored at cold temperatures (4□°C), as in Shiozaki et□al. (2021), is encouraged.

## DATA AVAILABILITY

Read counts following taxonomic assignment and bioinformatic filtering will be made available as a Supplementary file upon acceptance. Raw data (DNA sequences) are available in the European Nucleotide Archive (ENA) under the BioProject PRJEB97780 (https://www.ebi.ac.uk/ena/browser/view/ PRJEB97780).

## ACKNOWLEDGEMENTS

The Basque Government through its Consolidated Research Group “Genes, poblaciones, ecosistemas: investigación fundamental y aplicada (IT1571-22)” funded this research. We acknowledge the help of Natalia Miguens from SGIKER (UPV-EHU) and AZTI technicians in sample processing. Special thanks to Ann Bucklin (University of Connecticut), Rowena Stern (Marine Biological Association) and other ICES Working Group on Integrated Morphological and Molecular Taxonomy (WGIMT) colleagues for fruitful discussions.

