## Supplementary materials for "Is DNA metabarcoding an option for formaldehyde-preserved zooplankton time series?"

**-Supp. Mat. Table 1. Bioinformatic filtering details.** The number of sequences after filtering steps I and II (Figure 2) along with the copepods´ Shannon-Wiener H´ values are shown for the three extraction methods (HHA, HPC and SH), including the repaired (NextNEB FFPE repair kit) and non-repaired (original) sample pairs. A tick (√) signals those formaldehyde-fixed samples passing all the bioinformatic thresholds (see Methods for further information). Results for both the 18S V9 and the Günther miniCOI marker are shown (top and bottom tables, respectively). Samples´ CODEs as in Table 1.

**-Supp. Mat. Figure 1. Filtering of FF samples based on the number of reads and copepods´ Shannon-Wiener H´ values.** a) Copepods´ Shannon-Wiener H´ values for the copepod community based in microscopy and copepod biomass values; a threshold oh H´≥ 1.1 is defined (red line; see Methods for further information. b) Total number of 18S V9 sequences after bioinformatic filtering step-II (log_10_, bottom axis; Figure 2 and Supp. Mat. Table 1) plotted along with the H´ index value for the copepod 18S V9 sequences (left axis) for the formaldehyde-fixed samples (Supp. Mat. Table 1). Red lines signal the bioinformatic thresholds applied for further analysis.

**-Supp. Mat. Figure 2.** **DNA integrity (LabChip).** LabChip gel showing the degree of fragmentation for the DNA extracted from those 1995-2011 formaldehyde-fixed samples not shown in Figure 5. Samples´ CODEs in the bottom as in Table 1, extraction methods at the top of the image. Four negative controls (“BLANK”) are also shown.

**-Supp. Mat. Figure 3. Taxonomic composition of the positive controls (DNA metabarcoding).** Taxonomic characterization of the two positive controls (zooplankton samples fixed in ethanol, 22/C+1 and 22/C+2; Table 1 and Figure 1) for:a) 18S V9 b) Günther miniCOI and c) Meunier´s miniCOI. Samples´ CODEs as in Table 1. Taxa´s CODEs as in Table 2 for the 18S V9 marker.

**-Supp. Mat. Figure 4. PCR amplification gel for the Meusnier miniCOI marker (2017 and 2022 samples).** LabChip results for the Meusnier miniCOI marker (2017 and 2022 samples). The expected size of the band at this PCR is ~ 263 bp. Samples´ CODEs in the bottom as in Table 1, extraction methods at the top of the image. A positive control (22/C+1) is shown for comparison. Three negative controls (“BLANK”) are also shown.

**-DNA extraction and marker amplification protocols.** Detailed protocols for the three extraction methods tested here with formaldehyde-fixed samples (HHA, HPC and SH methods) along with the one applied to the positive controls (QIA; for ethanol fixed samples). DNA repair (NextNEB FFPE repair kit) protocol and PCR thermocycling details are also shown for the different markers tested.

**Supp. Mat. Table 1.**

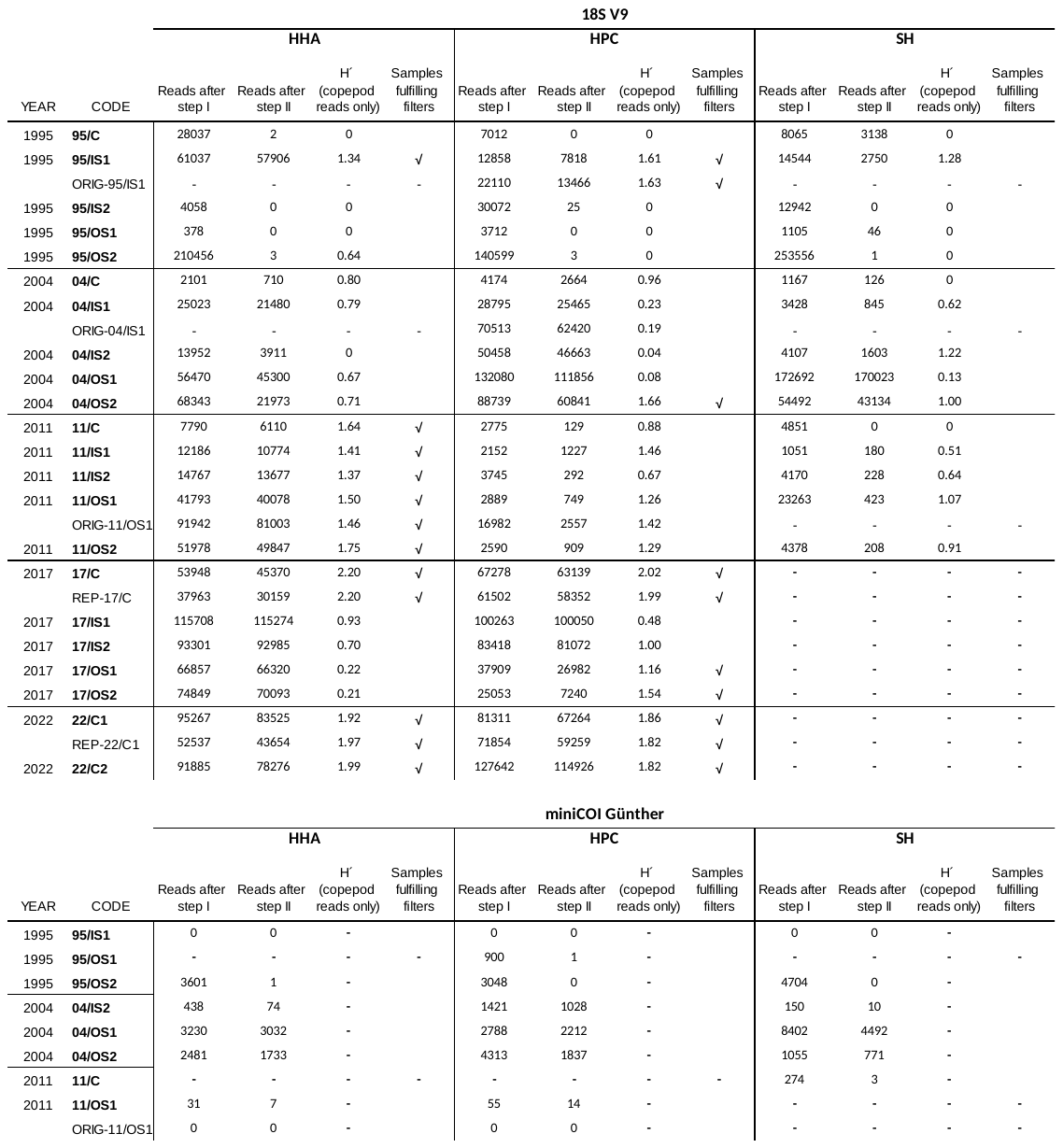

**Supp. Mat. Figure1.**

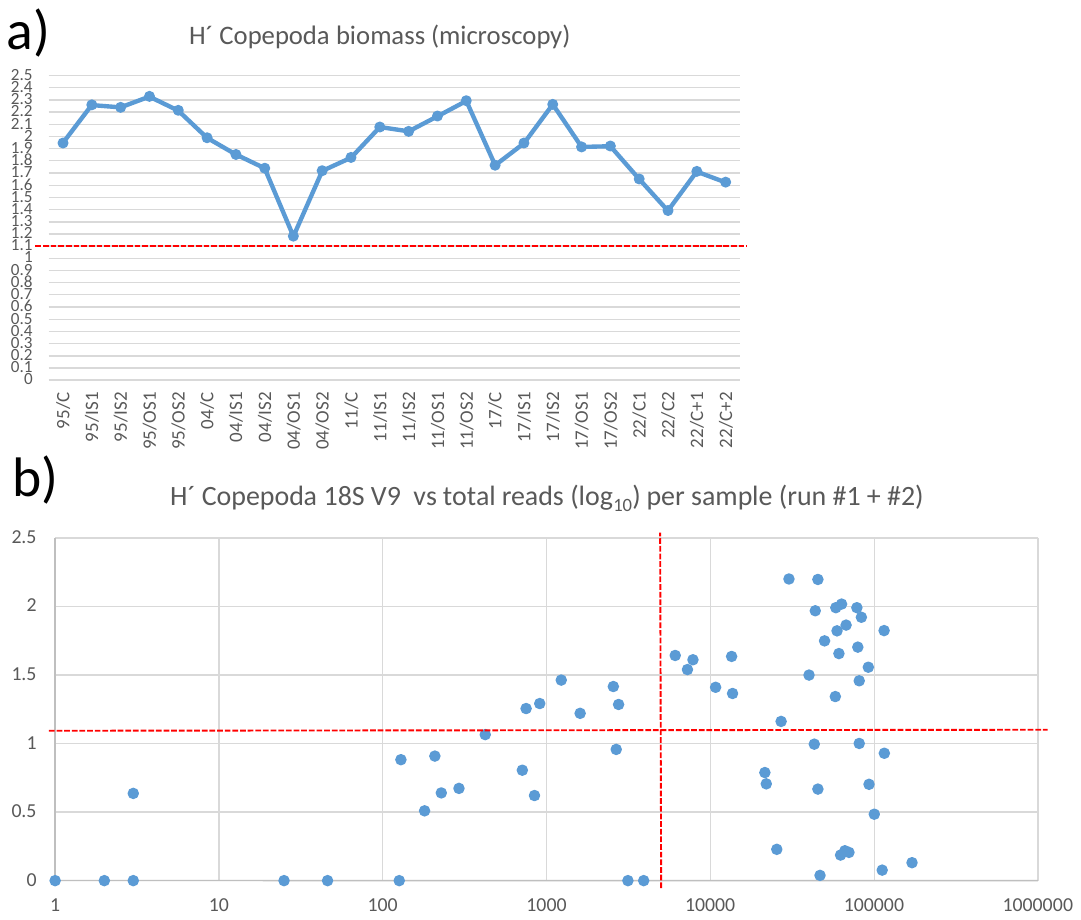

**Supp. Mat. Figure 2.**

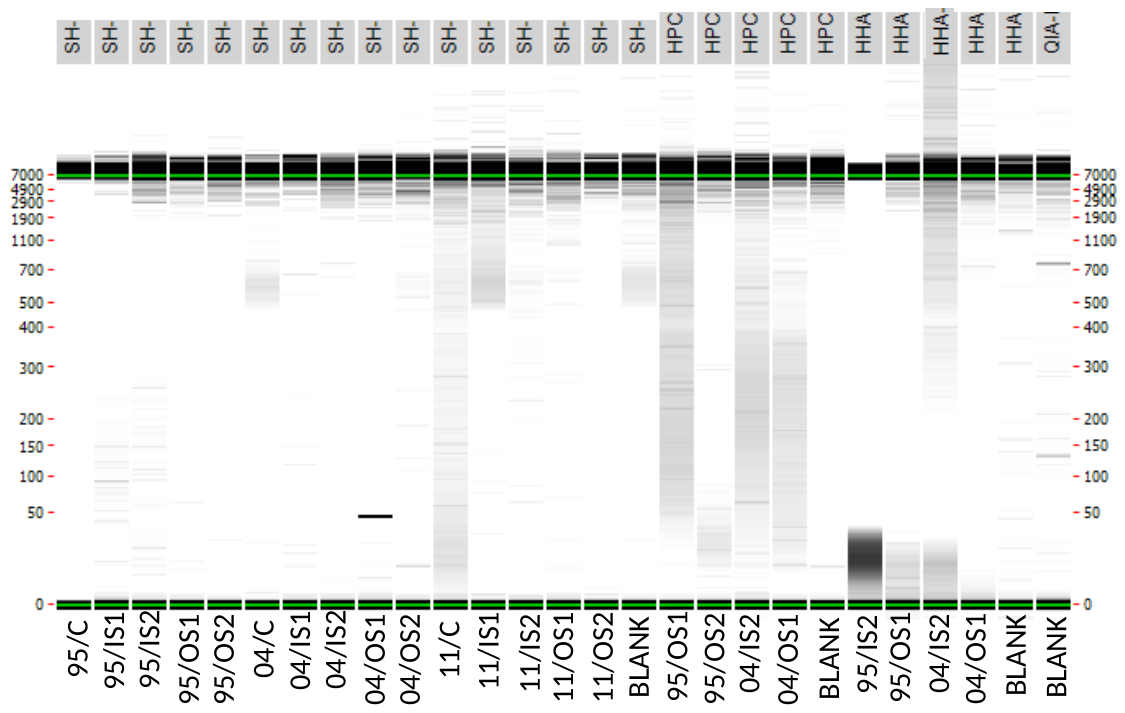

**Supp. Mat. Figure 3.**

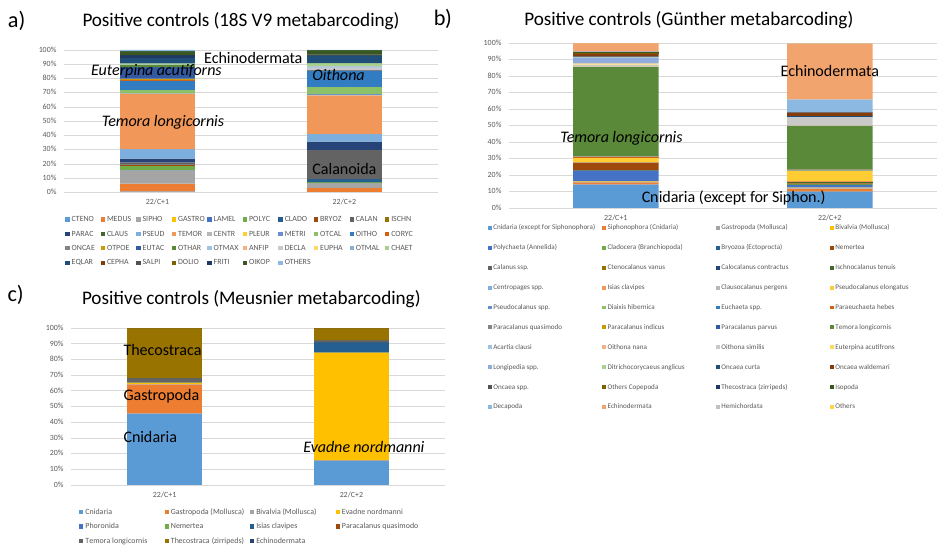

**Supp. Mat. Figure 4.**

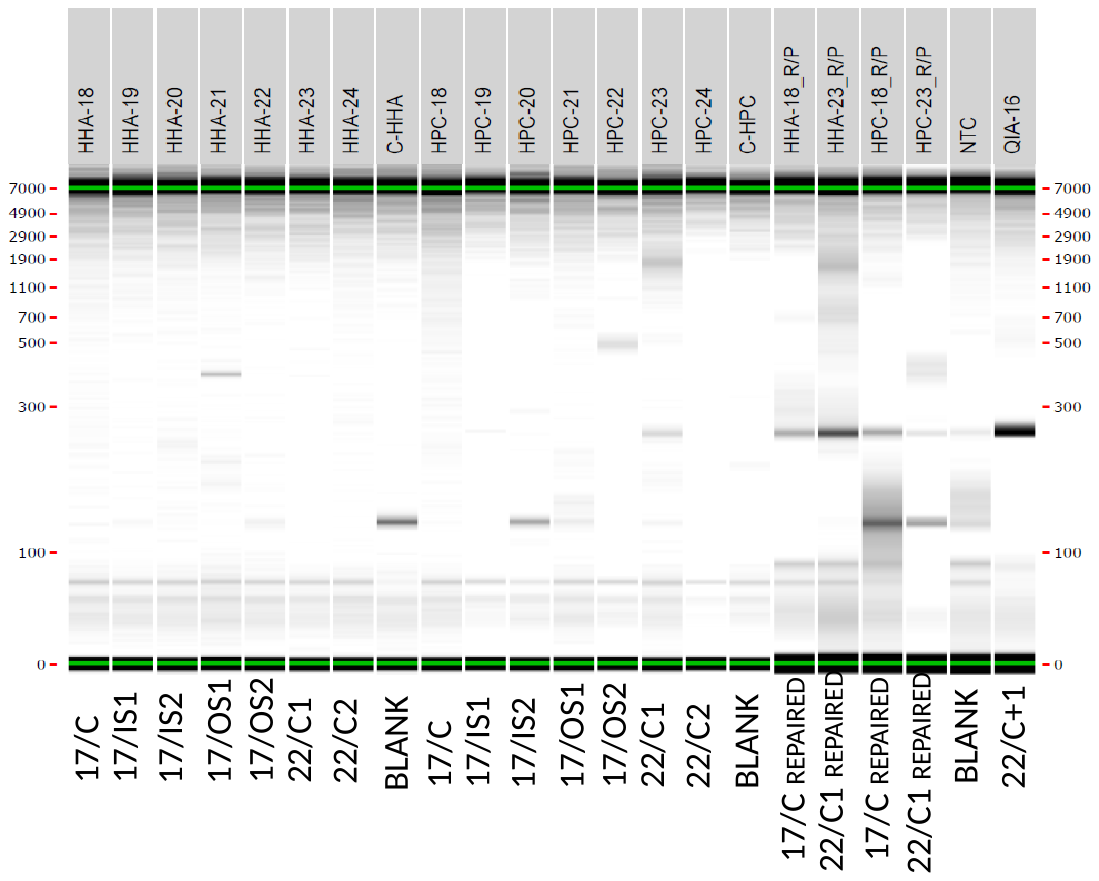

**DNA extraction and marker amplification protocols.**

**Previous steps - Formaldehyde-fixed samples´ preparation**

1. Centrifuge the 50 ml tubes at maximum speed for 10 min to obtain pellets.

2. Remove the ethanol supernatant without disturbing the pellets.

3. Weigh the tube relative to the empty tube to estimate the total mg of pellets.

4. Add up to 30 ml of cross-linked H2OmilliQ.

5. Vortex for 15 sec to wash the pellets.

6. Centrifuge the 50 ml tubes at maximum speed for 10 min to obtain pellets.

7. Remove the water supernatant without disturbing the pellets.

8. Repeat steps 4, 5, and 6 twice (three washes with cross-linked H2OmilliQ).

9. Collect up to a maximum of 250 mg in 2.0 ml homogenization tubes (avoiding collapse); three replicates as follows:

1. Original PowerSoil Pro tube, for Shiozaki (SH) extraction protocol.

2. Sarsted tube with a ceramic bead mix (glass powder to fill the conical section + 1.4 mm beads to 0.25 ml + 1 2.8 mm bead) for Hahn-Hot-Alkaline (HHA) extraction protocol.

3. Sarsted tube with a ceramic bead mix (glass powder to fill the conical section + 1.4 mm beads to 0.25 ml + 1 2.8 mm bead) for Hahn-Proteinase K-Column (HPC) extraction protocol.

10. Store frozen at -20 -30ºC until extraction.

**Buffer (lysis/digestion) composition**

-Shiozaki protocol (SH). BNB (20 ml): Borate-NaOH buffer with 1% SDS: 50 mM BNB (pH 11, adjusted with NaOH), SDS 1%:

• 2 ml of 0.5M Borate pH 8.5

• 1 ml of 20% SDS

• Bring to 20 ml with H2OmilliQ, adding 1M NaOH to pH=11 before finalising the volume.

-Hahn-Hot-Alkaline protocol (HHA). BA (12.5 ml): alkaline buffer (0.1 M NaOH with 1% SDS, pH 12-13):

• 1.25 ml of 1M NaOH

• 625 µl of 20% SDS

• Bring to 12.5 ml with H2OmilliQ, checking that the pH is between 12-13

-Hahn-Proteinase K-Column protocol (HPC). BL (20 ml): lysis buffer (10 mM NaCl, 20 mM Tris-HCl, pH 8.0, 1 mM EDTA, 1% SDS). The Tris-EDTA is 10 mM-1 mM.

• 200 µl of 1 M NaCl (2.92 g of NaCl dissolved in 50 ml of H2OmQ)

• 1 ml of 20% SDS

• Bring to 20 ml with 10 mM-1 mM Tris-EDTA

**Shiozaki protocol (SH; Shiozaki et al., 2021)**

1. Dry disruption of the tissue with Precellys Homogenizer (Bertin Technologies): 5000 rpm-2x12s-030s)-------spin.

2. Add 500 µl BNB (pH = 11) -------vortex 15”.

3. Wet disruption of the tissue with Precellys Homogenizer (Bertin Technologies): 5000 rpm-2x12s-030s)----spin.

4. Add 21.5 µl of pK at 18.6 mg/ml------vortex 15”

5. Digest for 24 hours at 65°C in a ThermoShaker (1200 rpm or maximum possible).

6. Continue the PowerSoil Pro (QIAGEN) protocol as follows

7. Centrifuge at 15,000 g for 1 min.

8. Transfer the supernatant to 2 ml tubes (usually 500-600 µl, but if there is more, recover all of it).

9. Add 200 µl of CD2 (stored at 4°C) to each tube.

10. Mix with a 5" vortex.

11. Centrifuge at 15,000 g for 1 min.

12. Transfer 700 µl of supernatant (do not collect particles) to a clean 2 ml tube.

13. Add 600 µl of CD3 to each tube.

14. Mix or vortex with a 5" vortex and spin.

15. Load the samples onto the filtration columns: 650 µl of the sample.

16. Centrifuge at 15,000 g for 1 min. Discard the filtrate.

17. Repeat this process until the entire sample volume has been filtered.

18. Place the filters in clean waste tubes.

19. Add 500 µl of EA solution to each tube.

20. Centrifuge at 15,000 g for 1 min. Discard the liquid.

21. Add 500 µl of C5 solution to each tube.

22. Centrifuge at 15,000 g for 1 min. Discard the liquid and place the filters in clean waste tubes.

23. Centrifuge at 16,000 g for 2 min.

24. Transfer the column to clean 1.5 ml tube.

25. Add 30 µl of C6 to each tube (Important: in the center of the membrane).

26. Let stand at room temperature for 2-3 min.

27. Centrifuge at 10,000 g for 1 min.

28. Remove the column.

29. Store the obtained DNA at -20 -30°C.

**Hahn-Hot-Alkaline protocol (HHA; adapted from Hahn et al., 2021, 2024 and Campos and Gilbert, 2012)**

1. Dry disruption of the tissue with Precellys Homogenizer (Bertin Technologies): 6500 rpm-3x30s-30s) --------spin.
2. Add 500 µl BA (pH=12-13)-------- vortex 15”.
3. Wet disruption of the tissue with Precellys Homogenizer (Bertin Technologies): 6500 rpm-3x30s-30s) ---- gentle spin.
4. Digest in a thermal block 100ºC for 40 min.
5. Transfer the supernatant to a clean 1,5 ml tube.
6. Allow the tissue-buffer to cool for 5 min to room temperature --------- spin.
7. Transfer 500 µl to a clean 1,5 ml tube.
8. Add 500 µl 25:24:1 phenol:chloroform:isoamyl alcohol to the mixture.
9. Mix gently at room temperature for 5 min.
10. Centrifuge for 5 min at >10,000 g to separate the layers.
11. Carefully remove the upper aqueous layer (be careful not to remove the protein-containing interface) and add to a new tube containing 500 µl chloroform. Discard the lower phenol layer properly.
12. Repeat steps 9–10.
13. Remove the upper aqueous layer and place in a new 1.5 ml eppendorf tube (estimate the collected volume). Discard the lower chloroform layer properly.
14. Add 1 volume isopropanol and 0.1 volume 3 M sodium acetate (approx. pH 5).
15. Immediately centrifuge at high speed (17,000 g) for 30 min at room temperature.
16. Immediately following centrifugation, decant the liquid from the tube carefully. The DNA will have precipitated into a pellet at the bottom of the tube and may not be visible.
17. To rinse the pellet, gently add 1 ml 85% ethanol, gently invert once, then centrifuge for 5 min at high speed (17,000 g).
18. Gently decant the ethanol.
19. Repeat steps 17-18.
20. All ethanol must be removed from the pellet as any residual ethanol will inhibit downstream applications. This can be easily achieved with a small bore pipette, followed by a brief incubation at a relatively high temperature (e.g., 60ºC for 5 min with the lids open).
21. Re-suspend the pellet in 30 μl TE - low EDTA. If the pellet has become very dry, leave it at room temperature in the liquid for 5–10 min, followed by gentle pipetting or vortexing.
22. Store the obtained DNA at -20 -30°C.

**Hahn-Proteinase K-Column protocol (HPC; adapted from Hahn et al., 2021)**

1. Dry disruption of the tissue with Precellys Homogenizer (Bertin Technologies): 6500 rpm-3x30s-30s) --------spin.
2. Add 500 µl BL (pH = 8)------vortex 15 sec.
3. Wet disruption of the tissue with Precellys Homogenizer (Bertin Technologies): 6500 rpm-3x30s-30s)-----spin
4. Add 32,2 µl pK (18,6 mg/ml) and digest in a ThermoShaker 65ºC for 24 h at 1200 rpm----spin
5. Add 5 volumes of Buffer PB to 1 volume of the sample, and then mix. The expected volume to use will be 300-350 µl. Thus:
   a. Transfer 325 µl to a clan 2,0 ml tube.
   b. Add 5*325 µl = 1625 µl PB.
   c. Add pH indicator (1:250, with respect to PB volumen): 6,5 µl pH Indicator.
6. Check that the mixture’s colour is yellow.
7. If the color of the mixture is orange or violet, add 10 µl of 3 M sodium acetate, pH 5.0, and mix. The color of the mixture will turn yellow.
8. Place a QIAquick spin column in a provided 2 ml collection tube.
9. To bind DNA, apply the sample to the QIAquick column and centrifuge 17000 g for 30–60 s.
10. Discard flow-through. Place the QIAquick column back into the same tube. Collection tubes are reused to reduce plastic waste. Repeat as many times as necessary until all the filtrate has passed through.
11. To wash, add 700 µl Buffer PE to the QIAquick column and centrifuge for 30–60 s (17000 g)
12. Discard flow-through and place the QIAquick column back into the same tube. Centrifuge the column for an additional 1 min. (17000 g).

IMPORTANT: Residual ethanol from Buffer PE will not be completely removed unless the flow-through is discarded before this additional centrifugation.

13. Place QIAquick column in a clean 1.5 ml microcentrifuge tube.
14. To elute DNA, add 30 ul buffer EB) to the center of the QIAquick membrane, let the column stand for 1 min and centrifuge the column for 1 min. (15000 g)
15. Store the obtained DNA at -20 -30°C.

**quiagen´s QIAamp DNA Mini Kit® (qia; adapted)**

1. Put 25-50 mg of fresh sample in a Sarsted type tube with a ceramic bead mix (glass powder to fill the conical section + 1.4 mm beads to 0.25 ml + 1 2.8 mm bead).
2. Dry disruption of the tissue with Precellys Homogenizer (Bertin Technologies): 6500 rpm-3x30s-30s). Gentle spin.
3. Add 100 µl PBS 1X ------ vortex 15”.
4. Wet disruption of the tissue with Precellys Homogenizer (Bertin Technologies): 6500 rpm-3x30s-30s). Gentle spin.
5. Add 180 ul Buffer ATL.
6. Add 20 ul pK.
7. Vortex and incubate in a ThermoShaker at 56ºC (1200 rpm) overnight.
8. Centrifuge the tube to remove drops from the inside of the lid.
9. Add 200 μl Buffer AL to the sample, mix by pulse-vortexing for 15 s, and incubate at 70°C for 10 min. Briefly centrifuge the microcentrifuge tube to remove drops from inside the lid.
10. Add 200 μl ethanol (96–100%) to the sample, and mix by pulse-vortexing for 15 s. After mixing, briefly centrifuge the microcentrifuge tube to remove drops from inside the lid.
11. Carefully apply the mixture to the QIAamp Mini spin column (in a 2 ml collection tube) without wetting the rim. Close the cap, and centrifuge at 6000 g (8000 rpm) for 1 min.
12. Place the QIAamp Mini spin column in a clean 2 ml collection tube (provided), and discard the tube containing the filtrate.
13. Carefully open the QIAamp Mini spin column and add 500 μl Buffer AW1 without wetting the rim. Close the cap, and centrifuge at 6000 g (8000 rpm) for 1 min.
14. Place the QIAamp Mini spin column in a clean 2 ml collection tube (provided), and discard the collection tube containing the filtrate.
15. Carefully open the QIAamp Mini spin column and add 500 μl Buffer AW2 without wetting the rim. Close the cap and centrifuge at full speed (20,000 g; 14,000 rpm) for 3 min.
16. Place the QIAamp Mini spin column in a clean 1.5 ml microcentrifuge tube (not provided), and discard the collection tube containing the filtrate.
17. Carefully open the QIAamp Mini spin column and add 55 μl H_2_O_milliQ_. Incubate at room temperature for 1 min, and then centrifuge at 6000 g (8000 rpm) for1 min.
18. Store the obtained DNA at -20 -30°C.

**DNA repair protocol**

A subset of samples was subjected to DNA repair using a commercial repair kit, the New England BioLabs' NEBNext FFPE DNA Repair v2 Module according to the manufacturer’s guidelines.

**Thermocycler programs**

- **18S V9**: an initial 15 min at 95ºC, followed by 30 cycles at 95ºC for 15s, 56ºC for 15s, 72ºC for 30s, and a final cycle of 7 min at 72 ºC.
- **Leray-Geller miniCOI**: an initial 95°C for 15 min, followed by a touchdown step of 10 cycles of 95°C for 10 s, 62°C (-1ºC each cycle) for 30s and 72°C for 60s, a second step of 25 cycles of 95°C for 10 s, 46°C for 30s and 72°C for 60s and a final cycle of 10 min at 72ºC (Leray et. al, 2013).
- **Meusnier miniCOI**: an initial 95°C for 15 min, 5 cycles of 95°C for 1min, 46°C for 1min and 72°C for 30s, a second step of 35 cycles of 95°C for 1min, 53°C for 1min and 72°C for 30s and a final cycle of 5 min at 72ºC.
- **Günther miniCOI:** an initial 95°C for 15 min, 35 cycles of 95°C for 30 s, 43°C for 30s and 72°C for 30s, and a final cycle of 7 min at 72ºC.
